# Bayesian generative modeling reveals a multi-modal hierarchical architecture in the mouse functional connectome

**DOI:** 10.64898/2026.06.01.729443

**Authors:** Srinivas Govinda Surampudi, Francesca Mandino, Evelyn Lake, Luiz Pessoa

## Abstract

Understanding the principles governing large-scale functional organization of the brain remains a central challenge in systems neuroscience. Despite convergent findings, substantial variability across analytical approaches suggests that functional networks may not admit a unique partitioning. Here, we propose that this variability reflects an intrinsic property of the connectome itself: its organization may be fundamentally multi-modal rather than singular. To test this hypothesis, we employ a Bayesian generative modeling framework based on stochastic block models, enabling principled comparison of competing organizational principles and characterization of the full posterior distribution over network partitions. Applying this framework to resting-state fMRI data in mice, we find that a non-degree-corrected hierarchical architecture provides the most parsimonious description of the functional connectome. Importantly, the inferred posterior landscape is not dominated by a single configuration, but instead comprises multiple distinct and co-dominant organizational schemes. At the mesoscale, these hierarchical communities are anatomically grounded yet systematically reorganize canonical resting-state networks: primary sensory systems remain cohesive, whereas higher-order association networks are fractionated into multiple interacting sub-circuits. This global structural variation is driven by structured variability at the community level, where integrative systems exhibit variable regional affiliations while sensory systems act as structurally stable anchors. Together, these findings suggest that the resting-state connectome is best described as a distribution over alternative, yet co-dominant, organizational configurations. This perspective reconciles inconsistencies across previous studies and supports a view of brain organization as inherently degenerate, providing a latent repertoire of network configurations that may underlie adaptive information routing and dynamic functional reconfiguration.

## 1 Introduction

Elucidating how distributed brain regions coordinate to form cohesive large-scale networks is a primary objective of systems neuroscience Sporns (2016); Bullmore and Sporns (2009); Bassett and Sporns (2017). While these macroscopic systems were originally mapped and extensively investigated in humans using task-free, resting-state functional MRI (rs-fMRI) Biswal et al. (1995); Fox and Raichle (2007); Damoiseaux et al. (2006), parallel research in rodents has become a vital vehicle for dissecting the core organizational principles of the connectome under highly controlled experimental conditions Gozzi and Schwarz (2016); Zerbi et al. (2015). Across both human and animal studies, multiple analytical techniques have been employed to define these networks, including Independent Component Analysis (ICA), modularity optimization, and co-activation patterns (CAPs), among others. Across studies, a few systems have been repeatedly observed, including visual, sensorimotor, and default-mode networks. However, considerable variability is observed across techniques and studies. We hypothesized that such variability reflects the fact that the organization of large-scale networks, as garnered via the correlation of time series data, is not necessarily unique (see Figure 1). In other words, whereas there may exist natural region clusters (e.g., sensorimotor regions), it is possible that the functional organization of the brain admits multiple credible (i.e., of similar “quality”) network-level partitionings Meunier et al. (2009); Hilgetag and Goulas (2020); Palla et al. (2005); Vafaii et al. (2024).

**Figure 1:**
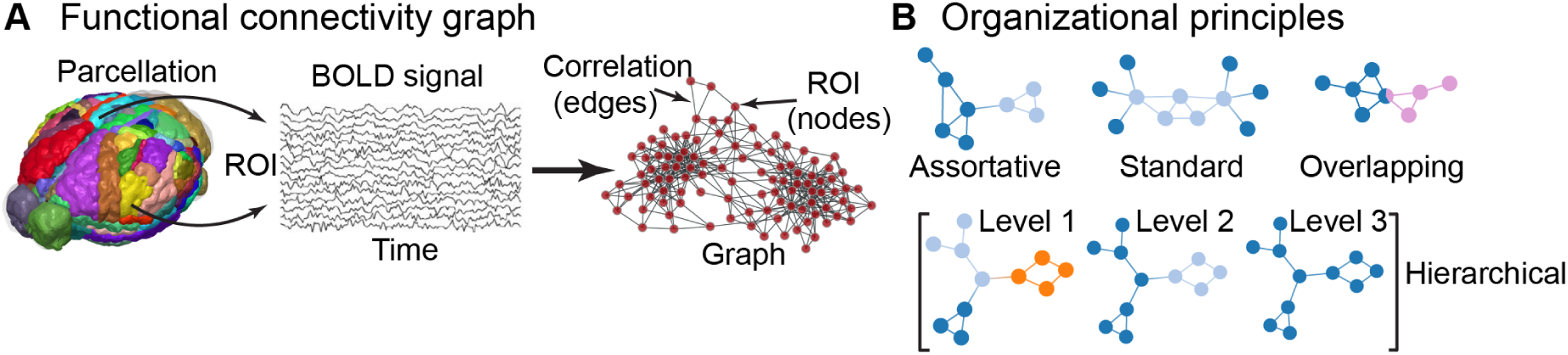
Functional brain networks can be described by multiple organizational principles. (A) Functional connectivity networks are constructed from fMRI data. Time series from brain regions of interest (ROIs) are correlated to create a graph where ROIs are nodes and strong correlations form the edges. (B) The resulting network can be organized according to several distinct principles, which represent competing hypotheses about its structure. These include purely “assortative” (segregated) communities, flexible “standard” communities that allow patterns like core-periphery, “overlapping” communities with mixed-membership nodes, and “hierarchical” arrangements with nested levels of organization.

Addressing this question is challenging because the different methods employed to investigate functional organization have distinct, and often implicit, technical properties. For example, a spatiotemporal technique, ICA Hyvärinen and Oja (2000), assumes that the underlying source signals (brain networks/components) are statistically independent of each other, such that activity in one network should not be predictable from the activity in another. Graph-based technique, modularity optimization methods Newman (2006); Fortunato (2010), maximize a global network partition quality often maximizing, in the context of fMRI data, “assortativity”, namely the tendency for nodes to be grouped with nodes with similar characteristics (e.g., activity). Whereas ICA networks can overlap spatially, modularity optimization algorithms assume non-overlapping (disjoint) organization (i.e., a region belongs to one and only one network). Another spatiotemporal technique, CAPs Liu and Duyn (2013); Liu et al. (2018), assume that brain activity can be characterized by a finite set of discrete, recurring spatial activation patterns or “states”. Unlike ICA’s continuous mixing in time, CAPs assume that at any given time, only one (or few) patterns are “active”, and that brain activity switches between distinct configurations.

It is not surprising, therefore, that many of the results obtained with these techniques differ in important ways, despite also capturing converging organization. Zerbi et al. (2015) applied group ICA Hyvärinen and Oja (2000) and identified 23 functional circuits hierarchically clustered into six bilaterally symmetric networks: somatosensory, sensory processing (motor, visual, and auditory), olfactory processing, limbic system (retrosplenial cortex, cingulate, hippocampus, thalamus, and amygdala), basal ganglia (caudate-putamen/dorsal-striatum, and pallidum), and cerebellar network. Grandjean et al. (2020) confirmed this organization across 17 independent center datasets using both group ICA and seed-based correlation analysis. Liska et al. (2015) used graph-theoretic modularity maximiza-tion Newman and Girvan (2004) and observed six communities, including default-mode (retrosplenial cortex, anterior cingulate, temporal cortex), lateral sensory-motor cortex, basal forebrain, ventral mid-brain (amygdala, hypothalamus), thalamus, and hippocampus networks. Coletta et al. (2020) using voxel-level modularity maximization, found four large-scale networks: default-mode plus limbic, lateral sensory-motor, hippocampal, and olfactory/basal forebrain/ventral midbrain networks. Finally, Mandino et al. (2022) using co-activation pattern (CAP) analysis Liu and Duyn (2013); Liu et al. (2018), discovered three networks, including default-mode and lateral sensory-motor cortex. Gutierrez-Barragan et al. (2019, 2022) investigated CAPs, and these preferentially highlight arousal-linked basal forebrain/thalamic engagement and anti-coordination between unimodal and polymodal cortices. In contrast, seed-based analysis in the same awake mice recovered default-mode, lateral cortical, and salience-like networks (similar to those found by Mandino et al. (2022)). For a detailed review of functional systems in rodents see Xu et al. (2022).

To investigate the potential multiplicity of network organizations from the same dataset, what is needed is a principled framework to evaluate the extent to which different organizational principles explain the fMRI data. Here, we employ a Bayesian generative modeling approach to do precisely that Peel et al. (2022); Peixoto (2019, 2023). Essentially, different techniques implicitly assume how the identified communities of brain regions interact with each other: assortative (disjoint partitioning), overlapping, and/or hierarchical arrangements (Figure 1). We make the assumptions, i.e. the organizational principles, explicit so that we can evaluate their compatibility with the data, thereby testing the presence of multiple organizations in the same dataset. The Bayesian framework then quantitatively evaluates compatibility between the organizational principles with the data on a single, information-theoretic scale, going beyond the technical differences between current methods. We chose to evaluate these organizations given their prevalence in the network neuroscience literature Sporns and Betzel (2016); Gozzi and Schwarz (2016); Najafi et al. (2016); Vafaii et al. (2024); Meunier et al. (2009); Betzel et al. (2024). It should be stated, however, that other forms of organizations have been reported in the literature, and that our results do not preclude that they are noteworthy organizational principles that explain some of the functional organization of brain large-scale networks; for example the framework of coactivation patterns discussed above.

Bayesian generative models Peixoto (2019); Peel et al. (2022) provide a principled framework for understanding complex systems by explicitly modeling how observed data arises from “network structure”, namely, an explicit probabilistic (mathematical) model of how the network is hypothesized to be constructed (Figure 2A). Stochastic block models (SBMs) Holland et al. (1983) are a powerful and versatile class of generative models that are capable of holding the organizational principles as their parameters, by virtue of which they generate the observed data (the functional connectivity network). The standard SBM Peixoto (2017) allows assortative, core-periphery, and bipartite organization patterns to emerge naturally Betzel et al. (2018b). A constrained variant explicitly generates purely assortative partitionings Zhang and Peixoto (2020), and specific extensions of the standard SBM allow overlapping Peixoto (2015) and hierarchical possibilities Peixoto (2014c). By fitting different stochastic block models (SBMs), we sought to quantitatively compare which organizational principle best explained the observed functional connectivity patterns through model selection based on posterior probabilities (see Figure 2B). We expressed the probabilities in terms of an interpretable quantity from information theory, called description length, that represents the explanatory efficiency of the organizational principle (see Section 4.3.1). In all, Bayesian generative models make the assumptions about data generation explicit and testable on an interpretable scale.

**Figure 2:**
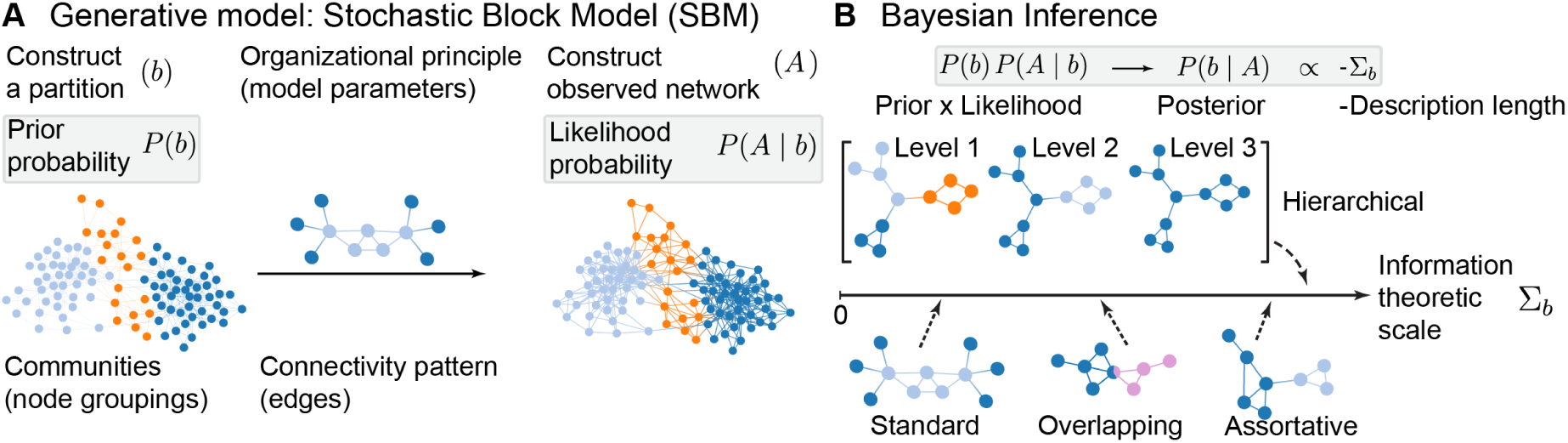
Bayesian generative framework allows for principled model comparison. (A) The generative model, here the stochastic block model (SBM), provides a formal recipe for network construction. It begins with a prior probability (P (b)) over possible community partitions. For a given partition, an organizational principle (e.g., assortative, hierarchical) defines a likelihood (P (A | b)) which is the probability of observing the network A given that partition structure. (B) Bayesian inference reverses this process. Given an observed network, it computes the posterior probability (P (b | A)), which quantifies how responsible the partition b is for generating the network A. The posterior is proportional to the likelihood times the prior. For model comparison, the posterior is expressed on a single information-theoretic scale using the description length (Σb)—number of bits required to describe the network in terms of that partition and the organizational principle; the lower the description length, the more efficient the model. The posterior is proportional to the negative of the description length; a model that provides a more efficient explanation (higher posterior, lower description length) is considered a better fit, allowing for a principled comparison between different organizational hypotheses.

Beyond comparing these broad organizational principles, a second challenge emerges. Traditional approaches to characterizing brain organization from fMRI data often seek a single partition that best summarizes functional organization, assuming that the space of solutions, or network partitions, concentrates around a single division of nodes into communities (a partition), with individual partitions being minor variations of their consensus at the center (Figure 3A; Lancichinetti and Fortunato (2012)). However, the pursuit of a consensus can be problematic when the underlying *distribution* of network partitions is inherently (or even moderately) heterogeneous Good et al. (2010); Fortunato (2010); that is, a single consensus can obscure the fact that the data may support multiple, distinct community arrangements. Figure 3B, for example, illustrates a potential case in which the landscape of solutions comprises of multiple concentrations of partitions, i.e. *solution modes*, such that partitions within a single mode are minor variations of their local consensus, but partitions in different modes are “structurally” distinct. In the example, we observe not a single dominant organization but rather multiple distinct modes that coexist (in this case, five), where a single consensus solution is unable to represent either of the distinct solutions. Existence of multiple modes in the solution landscape means that brain’s functional organization may not be a single, fixed map of community assignments, but rather be described as a landscape of coexisting distinct arrangements. We hypothesize, therefore, that the brain’s functional architecture may be partitioned in multiple distinct ways, even under the same organizational principle. Each mode represents a different possible arrangement of communities, potentially reflecting different complementary aspects (states, individual variations) of functional architecture that cannot be summarized by a single partition alone Mattar et al. (2015); Betzel et al. (2017).

**Figure 3:**
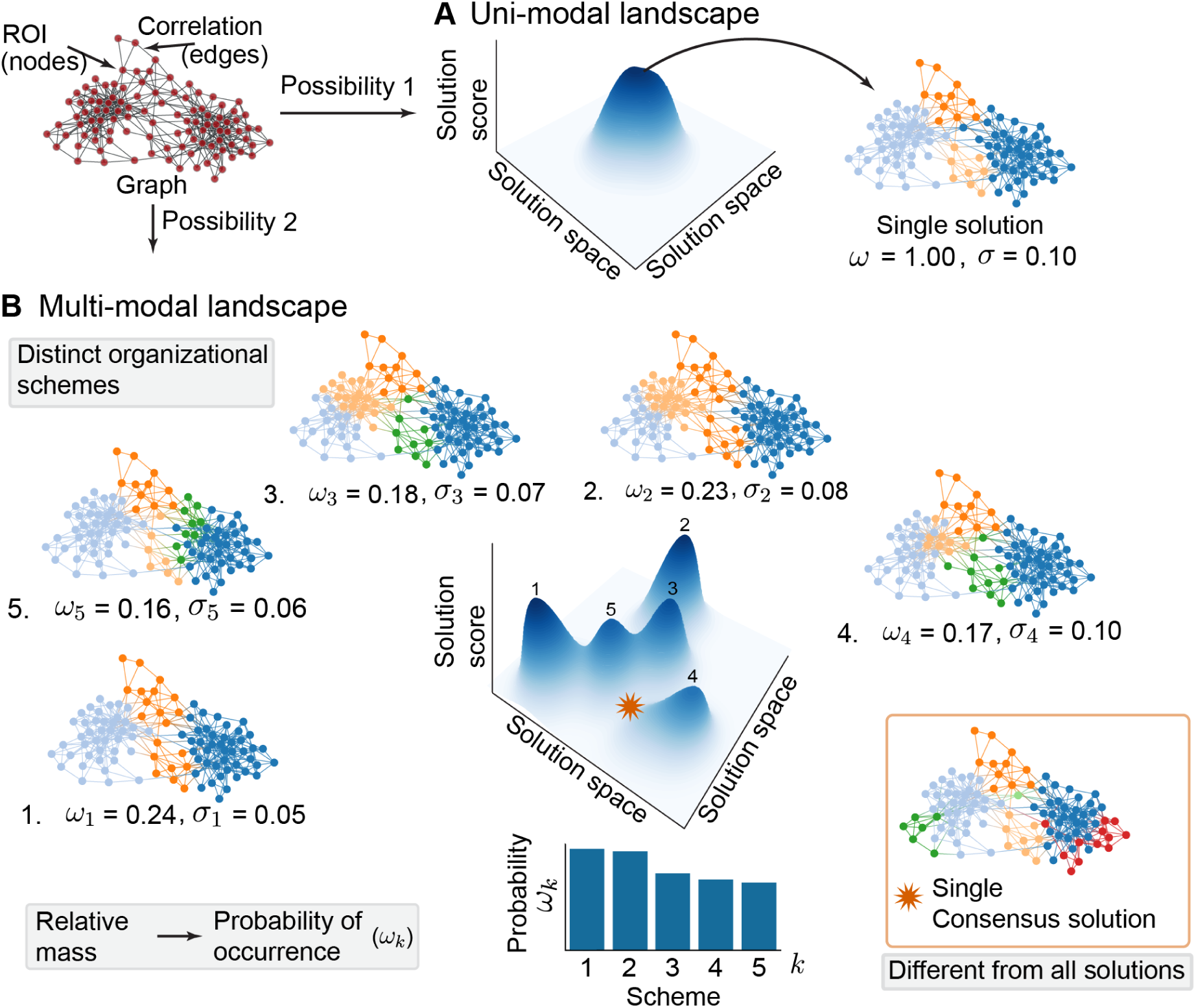
Brain network organization may reflect a multi-modal landscape of solutions. (A) A unimodal landscape represents the traditional assumption that a network’s organization can be accurately summarized by a single consensus partition. In this scenario, all possible solutions are minor variations of one central structure. (B) A multi-modal landscape reflects the hypothesis that the data may support multiple, structurally distinct organizational schemes, or solution modes. Each model represents a different, coherent way to partition the network. Critically, a single consensus partition (star) can be a poor summary, potentially corresponding to a low-probability solution that does not resemble any of the valid schemes. The probability of each scheme is quantified by its weight (*ω_k_*), denoting its prevalence in the landscape, and the variability within a scheme is captured by its uncertainty (*σ_k_*).

The Bayesian methodology we employ is uniquely suited to discover multiplicity of arrangements within an organizational principle Peixoto (2021). By characterizing the full posterior distribution over all possible partitions, it allows us to identify multiple solution modes if the data support their existence. Furthermore, it assigns a relative weight, or probability (*ω*), to each mode, quantifying its prevalence in the solution landscape (see Figure 3B). The fact that multiple modes can have substantial weight means the data support multiple, potentially complementary ways of understanding the brain’s functional architecture, rather than there being a single “correct” partition.

## 2 Results

To investigate the large-scale functional organization of the mouse brain, we analyzed resting-state BOLD-fMRI data from lightly anesthetized mice (*n* = 9, 0.5 − 0.75% isoflurane). Data were collected across three longitudinal sessions, each comprising four runs that lasted 10 min long, yielding a total of 120 min of data per mouse (see Section 4.1 for details). Following standardized preprocessing Desrosiers-Gŕegoire et al. (2024) and co-registration to the Allen CCFv3 space Wang et al. (2020), we constructed group-level functional connectivity graphs comprising of *N* = 172 brain regions of interest (ROIs) (see Section 4.1.5 for details). We defined each ROI by merging the regions in the CCFv3 based on their functional roles and anatomical proximity.

To construct a sparse, undirected graph suitable for network analysis, the group-level Pearson correlation matrices were binarized by retaining only the top *d*% strongest edges (main text reports *d* = 20%; see Supplement for *d* = 10%). This standard thresholding step is necessary to remove spurious, low-magnitude noise correlation, thereby reducing computational burden and helping the generative algorithms isolate the true topological structure of the network.

To ensure our findings are robust to sampling variability, we embedded this network construction within a bootstrap resampling framework (see Section 4.1.6 for details). In addition to constructing a graph from the original cohort of mice (herein termed the *reference sample*), we generated 500 independent group-level graphs by sampling the subjects with replacement. This allowed us to provide confidence intervals and statistical testing for all subsequent metrics.

### 2.1 Hierarchical organization is more compatible with the resting-state functional organization than purely assortative organization

To evaluate the extent to which different organizational principles explain the resting-state functional organization in mice, we employed a Bayesian generative modeling framework Peel et al. (2022) (see Section 4.2 for details). This framework involves defining a “generative model” (that is, a probabilistic recipe) to construct the observed functional graph in terms of arrangements of latent communities (“partitions” of the graph). Crucially, these arrangements are dictated by a specific organizational principle (such as strict modularity or a nested hierarchy). A Bayesian inference procedure then inverts this generative process to infer the community structures responsible for the observed graph, calculating the posterior probability distribution over all possible community arrangements allowed by that specific rule.

We compared seven generative hypotheses (i.e., organizational principles) implemented as stochastic block model (SBM) variants Peixoto (2019). To formalize the most prominent topological theories in network neuroscience into testable models, we evaluated a purely assortative architecture Sporns and Betzel (2016) alongside standard Betzel et al. (2018b), hierarchical Hilgetag and Goulas (2020), and overlapping Najafi et al. (2016); Vafaii et al. (2024) arrangements. Furthermore, the standard, hierarchical, and overlapping models were parameterized both with and without degree-correction (see Section 4.2.2 for details).

This distinction between non-degree-corrected and degree-corrected models allowed us to test a fundamental assumption regarding how highly connected regions (“hubs”) are integrated within the network. Specifically, this distinction evaluates topologies often associated with “rich-clubs” — densely interconnected communities of hubs that form a centralized communication backbone Sporns and Betzel (2016); Sporns (2016); Bullmore and Sporns (2009). Non-degree-correction assumes that a region’s connectivity is a homogeneous feature of its community, thereby grouping highly connected nodes together in a manner that can capture rich-club-like topographies. Conversely, degree-correction allows nodes with widely varying connectivity profiles to be grouped together Karrer and Newman (2011), implying that hubs are distributed across heterogeneous functional systems rather than forming an exclusive core. Finally, because modularity maximization is commonly used to investigate large-scale networks Liska et al. (2015); Coletta et al. (2020), we included the purely assortative SBM as its generative equivalent (see Section 4.2.3).

For each generative model, we approximated the posterior distribution over all possible graph partitions using Markov chain Monte Carlo (MCMC) simulations (see Section 4.2.4 for details). Briefly, the algorithm systematically navigates the vast landscape of possible community arrangements (partitions) to isolate the specific ensemble of partitions most strongly supported by the data. To objectively adjudicate between these competing hypotheses, we quantified each model’s “explanatory power” using Total Description Length (TDL; see Section 4.3.1 for details). Grounded in information theory, TDL represents the total number of bits required to succinctly describe the network data under a given generative rule. TDL balances goodness-of-fit against the complexity of the solution landscape. Models with a lower TDL encode the data more efficiently, making them more compatible with the network’s underlying topology.

Model comparison based on TDL revealed a clear and consistent ranking of organizational principles (Figure 4). The non-degree-corrected hierarchical model (nd-h-SBM) yielded the lowest TDL, indicating that a degree-homogeneous nested (multi-scale) architecture is more compatible with the underlying functional connectome of the mouse brain than the other evaluated principles. Conversely, the purely assortative model (a-SBM) — the generative equivalent of strict modularity maximization — yielded the highest description length. This result suggests that characterizing the brain solely as a system of strictly segregated modules provides a less compatible description of the network’s architecture, demonstrating that pure assortativity alone may be insufficient to capture the full complexity of the functional connectome.

**Figure 4:**
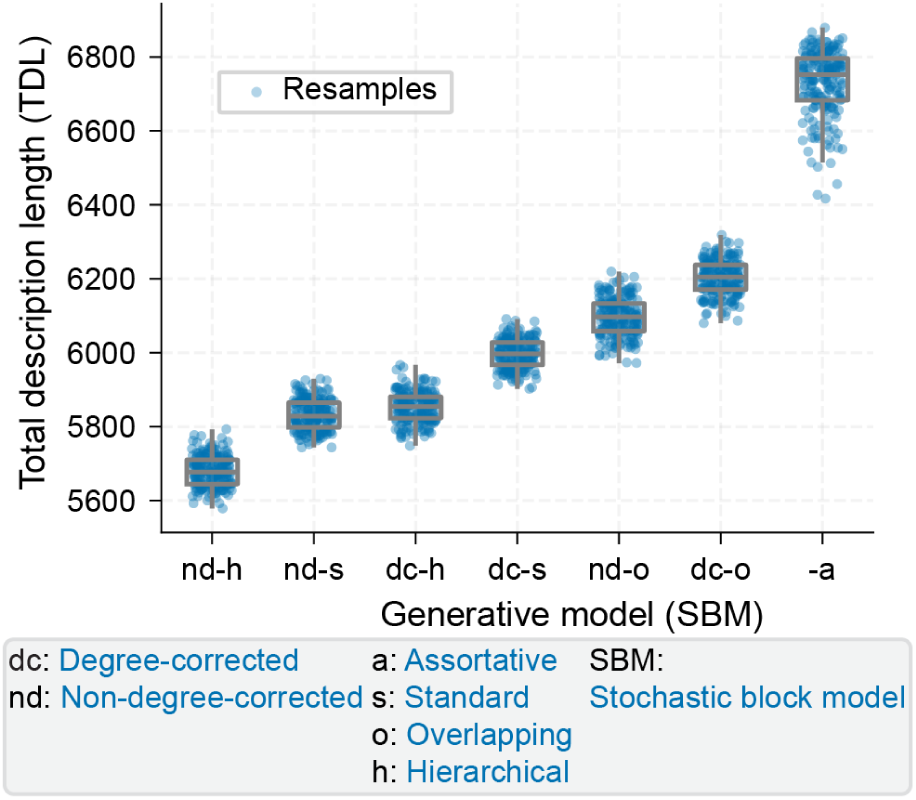
Hierarchical, degree-homogeneous communities are highly compatible with the functional connectome. Model comparison based on Total Description Length (TDL) across seven competing generative hypotheses. Box plots summarize the distribution of TDL values across 500 bootstrap resamples. Lower TDL indicates a more parsimonious and compatible explanation of the functional graph. The non-degree-corrected hierarchical model (nd-h-SBM) consistently yielded the lowest TDL, while the purely assortative model (a-SBM) — the generative equivalent of modularity maximization — yielded the highest. Across all tested organizational principles, the non-degree-corrected models (nd) were consistently more compatible than their degree-corrected (dc) variants.

Furthermore, across all evaluated hypotheses, non-degree-corrected models were consistently more compatible with the data than their degree-corrected variants. This structural preference indicates that, within this generative framework, high connectivity is modeled as a collective property shared within specific communities rather than an independent trait of individual regions. While this topological pattern resembles “rich-club” organizations (densely interconnected groups of regions that jointly facilitate global communication Bullmore and Sporns (2009); Sporns and Betzel (2016)) this preference may also be influenced by the inherent degree homogeneity of the thresholded BOLD network (discussed further in Section 3). Importantly, the overall ranking of these organizational principles remained robust to variations in edge density thresholding (see Supplement 5.5.1).

### 2.2 The inferred communities are anatomically grounded

Having identified the non-degree-corrected hierarchical principle as the most compatible generative rule, we first sought to contextualize its inferred communities before exploring the multiplicity of the solution landscape. As a reminder, while the network nodes were defined using the Allen CCFv3 atlas, the Bayesian generative models were not provided with any explicit spatial priors regarding how these regions should be grouped into communities. It is therefore critical to determine whether these algorithmically derived communities correspond to anatomically meaningful systems or are merely mathematical artifacts.

To extract these communities, we analyzed the full solution landscape of graph partitions generated by the MCMC simulations (see Section 4.2.4). The simulations approximate the posterior landscape by collecting a vast ensemble of valid partitions, where each partition represents a discrete assignment of ROIs into communities governed by the generative rule (here, hierarchical). To synthesize this massive ensemble and account for potential multi-modality in the landscape, we applied a mixture model Peixoto (2021) that grouped similar partitions into distinct clusters. Each cluster represents a distinct *organizational scheme*, and the relative size of the cluster reflects that scheme’s prevalence, or probability (*ω*), within the landscape (see Section 4.3.2 for details). After computationally aligning the partitions to ensure consistent community identities across the ensemble (and bootstrap resamples), we summarized the ROI community assignments within each scheme as a probabilistic assignment matrix (*π*) by aggregating over the partitions within that scheme. In this matrix, each column represents a community, and the values quantify the probability that a given ROI is assigned to that community within that specific scheme.

To identify the overall ROI assignments to the communities across this multi-modal landscape, we computed the expected assignment probability of each ROI by taking a weighted average (using *ω*) of the *π* matrices across all schemes. To evaluate which ROIs are assigned to which communities, we used a statistical test across the 500 bootstrap resamples to separate robust topological structure from background noise. We retained only those ROIs whose expected assignment probability was statistically greater than zero (FDR-corrected, *α* = 0.05; see Section 4.3.3). This filtering procedure isolated the highly reliable member ROIs of each community.

Because the hierarchical model was identified as the most compatible generative rule, it yielded a distinct probabilistic assignment matrix for each hierarchical level. Briefly, the ground level yielded over 20 spatially compact communities (Figure S11, S12). These merge to form 9 communities at the middle level (Figure 5), which further consolidate into broad macro-systems at the upper level, largely segregating the massive cortico-striatal forebrain from the brainstem and hindbrain (Figure S13). Here, we focus on the communities at the middle level due to their spatial comparability to canonical resting-state networks (RSNs) (Figure 5). Notably, despite the generative model receiving no explicit spatial priors during inference, the inferred communities emerged as anatomically contiguous and bilaterally symmetric. We assigned anatomical names to each community based on their constituent regions. Crucially, the neurobiological validity of these functional axes is not limited to prior resting-state fMRI literature, but is also grounded in multimodal metrics, including viral axonal tracing Oh et al. (2014); Zingg et al. (2014) and behavioral lesion studies Cardinal et al. (2002); Saper et al. (2001).

**Figure 5:**
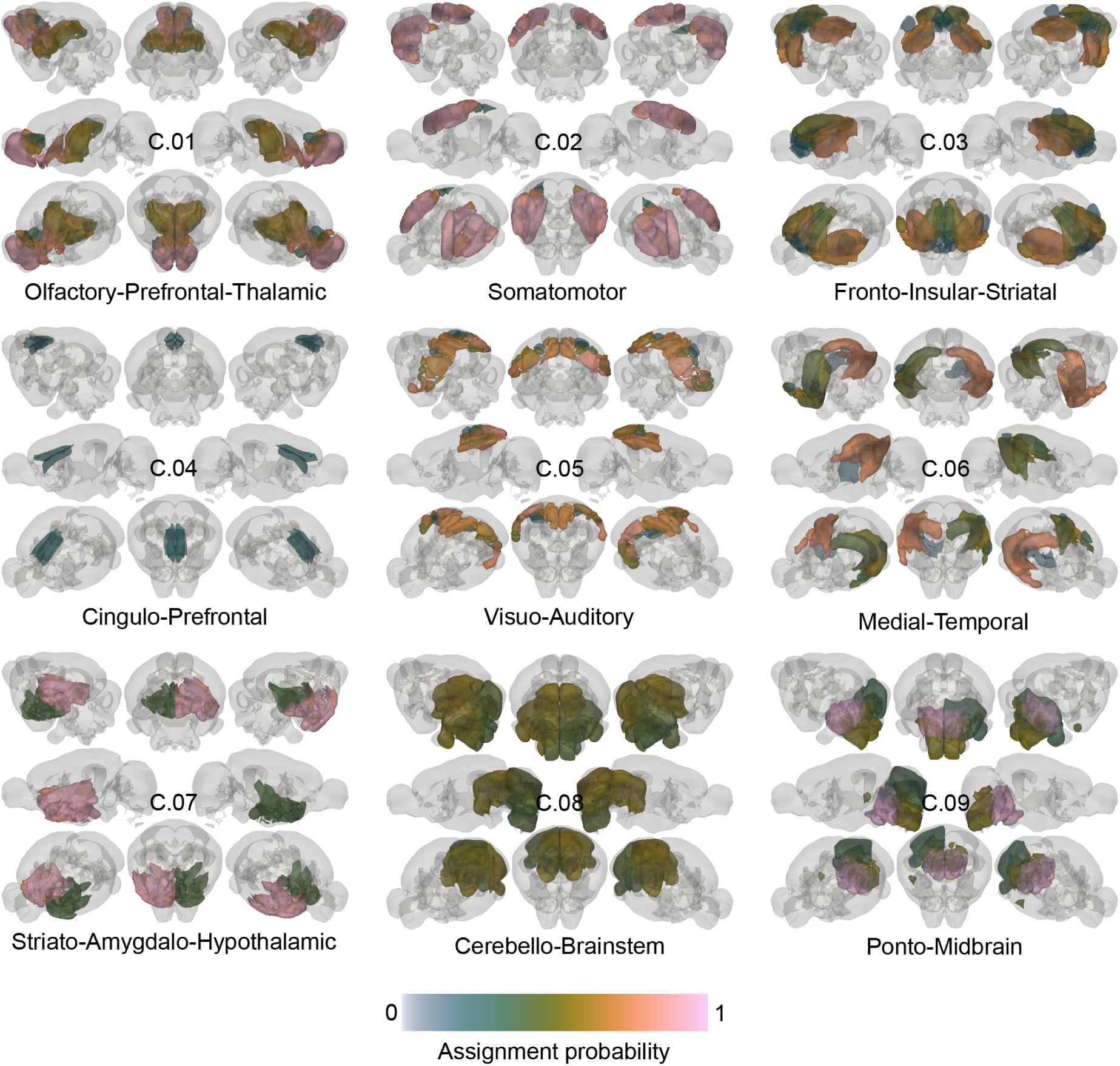
The most compatible generative rule discovers anatomically meaningful functional units of the mouse brain. Expected ROI compositions of the nine middle-level communities inferred by the non-degree-corrected hierarchical model (nd-h-SBM). Brain maps display the expected membership probabilities of ROIs, computed by averaging across the multi-modal solution landscape. Only highly reliable constituent ROIs (expected membership probability statistically greater than zero across 500 bootstrap resamples; FDR-corrected, *α* = 0.05) are visualized; color intensities reflect the median membership probability across these resamples. Despite receiving no spatial priors during inference, the generative framework partitions the functional connectome into bilaterally symmetric and contiguous communities that strongly align with known structural pathways and broad functional axes.

Within the cortex, the model delineated the Somatomotor community (C.02), encompassing primary somatosensory and motor cortices — regions united by dense local structural connectivity and jointly recruited during sensorimotor behavior Oh et al. (2014); Gozzi and Schwarz (2016). Posteriorly, it identified the Visuo-Auditory community (C.05), grouping primary visual, auditory, and retrosplenial areas, which share reciprocal axonal projections supporting multisensory integration Oh et al. (2014). Anteriorly, the model identified two distinct prefrontal systems: the Cingulo-Prefrontal community (C.04), anchored by anterior cingulate areas frequently implicated in executive control, and the more distributed Olfactory-Prefrontal-Thalamic community (C.01). This latter community links the olfactory bulb to medial prefrontal, septal, and thalamic structures, reflecting known anatomical pathways associated with odor-driven associative learning Zingg et al. (2014); Oh et al. (2014).

Subcortical and limbic structures segregated into highly precise anatomical units. The Medial-Temporal community (C.06) isolated the hippocampus and parahippocampal cortices, a densely inter-connected network classically associated with spatial navigation and memory. Affective, autonomic, and basal ganglia circuits were partitioned into two distinct cortico-subcortical loops: the Fronto-Insular-Striatal community (C.03), anchoring the dorsal striatum and insula (regions linked via known striatal projection pathways and often coactivated during interoception), and the Striato-Amygdalo-Hypothalamic community (C.07), capturing the amygdala, ventral striatum, and hypothalamus — a structurally coupled axis widely implicated in affective processing and survival drives Cardinal et al. (2002).

Finally, deep brainstem and hindbrain regions were distinctly partitioned, reflecting their distinct projection targets and physiological roles. The model grouped the cerebellum, medulla, and sensorimotor midbrain into the Cerebello-Brainstem community (C.08), reflecting pathways supporting motor coordination, while isolating the pons and behavioral state-related midbrain into the Ponto-Midbrain community (C.09), a structural hub associated with basic arousal and physiological state regulation Saper et al. (2001).

Importantly, the emergence of these anatomically meaningful boundaries was highly robust to variations in edge density thresholding, yielding a consistent organization consisting of seven consolidated communities at a 10% threshold (see Supplement 5.5.2). The emergence of these anatomically coherent communities, derived entirely without explicit spatial priors during inference, lends strong credibility to the utility of our Bayesian generative modeling framework for mapping the functional connectome.

### 2.3 The inferred communities refine canonical resting-state networks

We next investigated whether the inferred communities directly align with canonical functional systems, or if they structurally refine them into distinct sub-circuits. To provide a reference frame for this comparison, we utilized the resting-state network (RSN) atlas from Zerbi et al. (2015), which was validated across 17 independent datasets Grandjean et al. (2020), establishing it as a robust standard in the field (Figure 6A). This atlas partitions the brain into six functional systems: the somatosensory (anchored by primary motor and somatosensory cortices), sensory (visual and auditory cortices), olfactory (olfactory bulb and prefrontal areas), limbic (hippocampus, amygdala, thalamus, and cingulate), basal ganglia (striatum), and cerebellar networks.

**Figure 6:**
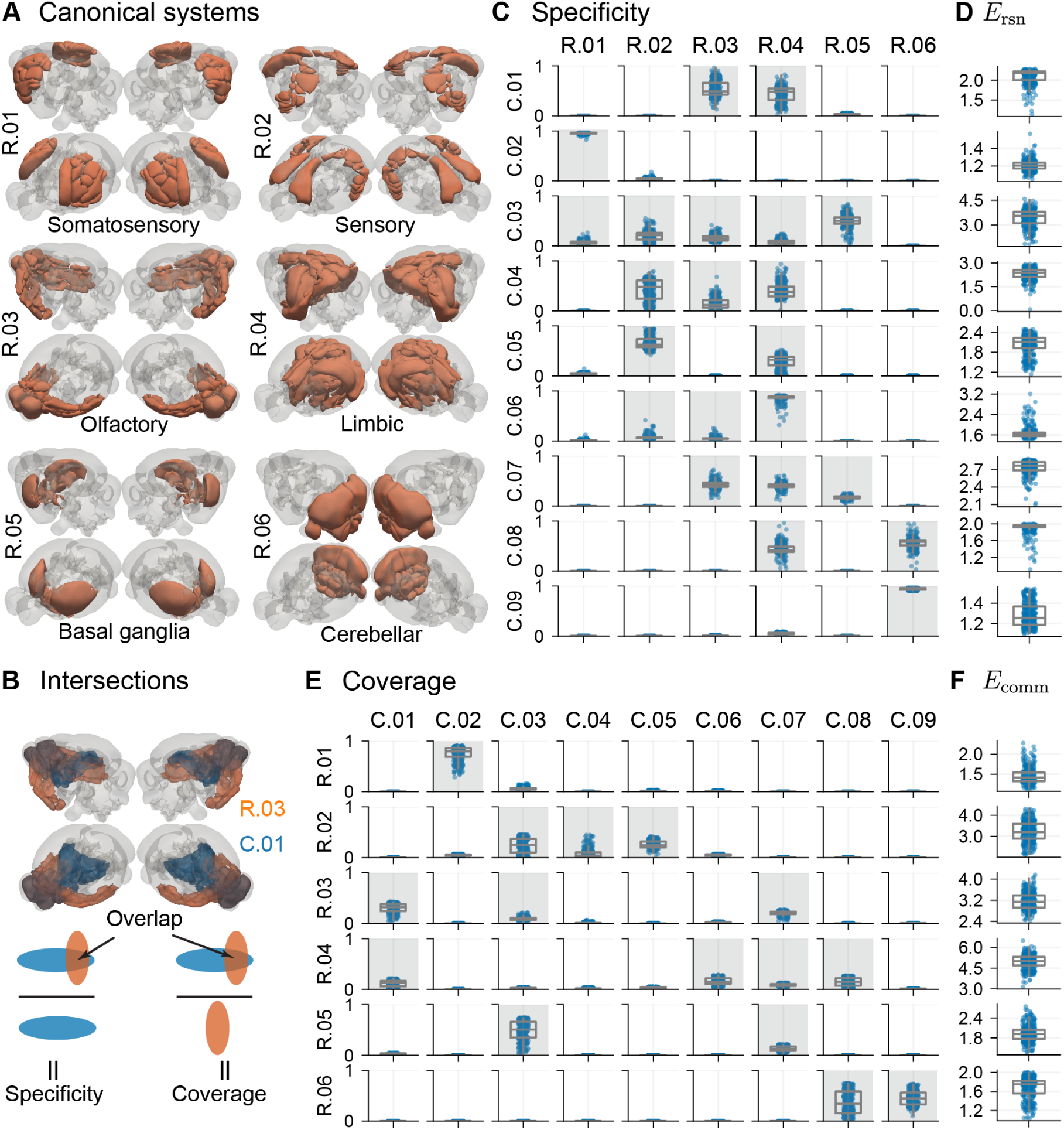
Inferred communities refine the boundaries of canonical resting-state networks. (A) The six canonical resting-state networks (RSNs) used as a reference atlas Zerbi, et al. (2015). (B) A schematic illustrating the bidirectional geometric metrics. Specificity is the fraction of an inferred community’s volume that falls within a given RSN. Coverage is the fraction of an RSN’s volume captured by a given community. In this example, the intersection is illustrated using the olfactory bulb. (C, D) From the community’s perspective, specificity profiles (C) are summarized by the effective number of RSNs spanned (*E*_rsn_) (D), yielding a measure of integration. (E, F) From the RSN’s perspective, coverage profiles (E) are summarized by the effective number of communities covering an RSN (*E*_comm_) (F), a measure of RSN cohesion vs. structural subdivision. In all box plots, each data point represents a single bootstrap resample (out of 500 total resamples). To generate each point, the metrics were first computed for each organizational scheme within that resample and then combined via a weighted average using the scheme probabilities (*ω*). The resulting distributions thus account for the multi-modal nature of the solution landscape.

To quantify the structural correspondence between our inferred communities and these canonical systems, we performed a bidirectional geometric analysis (Figure 6B; see Section 4.3.4 for details).

First, from the community’s perspective, we calculated *specificity* — the fraction of a community’s volume that intersects with a given RSN. Second, from the RSN’s perspective, we calculated *coverage*—the fraction of an RSN’s volume captured by a given community. Crucially, to ensure a reliable comparison, our inferred communities were strictly defined by their reliable constituent ROIs (Figure 5). Consequently, the total coverage of an RSN may be less than 100%.

Examining this geometric mapping from the perspective of the inferred communities (specificity) revealed a spectrum of structural integration (Figure 6C). To concisely summarize each community’s specificity profile across the canonical atlas, we computed the effective number of spanned RSNs (*E*_rsn_) using Shannon entropy (see Section 4.3.4 for details; essentially, a community entirely contained within a single RSN yields *E*_rsn_ = 1, whereas a community distributed perfectly evenly across two RSNs yields *E*_rsn_ = 2). While a minority of communities were highly localized within single canonical systems (*E*_rsn_ ≈ 1), the majority spanned multiple RSNs, potentially acting as integrative units (Figure 6D). For instance, the Somatomotor community (C.02) was highly specific, localizing almost entirely (95.4% median overlap across 500 bootstrap resamples) within the canonical somatosensory network (R.01), yielding a highly segregated *E*_rsn_ of 1.2. Conversely, the Fronto-Insular-Striatal community (C.03) exhibited a highly distributed specificity profile, spreading its volume primarily across the canonical basal ganglia (50.1%), sensory (21.5%), and olfactory (14.6%) systems. This wide distribution translates to an effective span of *E*_rsn_ = 3.55, quantitatively capturing its potential role as a bridge across multiple canonical domains.

Conversely, viewing the relationship from the perspective of the canonical systems (coverage) revealed how these traditional functional networks are composed (Figure 6E, F). By calculating the effective number of covering communities (*E*_comm_), we observed a clear spectrum from cohesion to structural subdivision. Primary sensory systems remained largely cohesive; for example, the canonical somatosensory network (R.01) was structurally unified, heavily captured by the single Somatomotor community (C.02; 79.5% coverage; *E*_comm_ = 1.41). In stark contrast, higher-order association networks were heavily subdivided. Most notably, the canonical limbic system (R.04) lacked a singular structural backbone (*E*_comm_ = 5.01). This broad functional territory was structurally distributed, with its volume divided primarily across the Olfactory-Prefrontal (C.01; 12.3%), Medial-Temporal (C.06; 14.9%), and Striato-Amygdalo-Hypothalamic (C.07; 7.9%) communities, alongside hindbrain contributions.

Together, these bidirectional metrics demonstrate that our generative framework refines the canonical functional atlas in a principled manner. The high specificity (segregation) and coverage (cohesion) of primary sensory systems suggest that, at the periphery, functional synchronization maps cleanly onto localized structural wiring. However, in higher-order cortex and subcortex, this 1-to-1 mapping dissolves. The generative model reveals that broad networks (such as the canonical limbic system) are actually composite structures formed by multiple distinct sub-circuits. Reciprocally, the inferred communities routinely span across these traditional RSN boundaries, structurally integrating them in a manner that potentially supports complex multi-system behavior.

### 2.4 The resting-state functional connectome is supported by multiple co-dominant organizational schemes

In the preceding analyses, we investigated the expected ROI assignments to the inferred communities by aggregating across the solution landscape (the posterior distribution over all possible partitions). Here, we investigated whether this landscape is characterized by a singular dominant organization, or if it is inherently multi-modal, sustaining multiple distinct but equally valid organizational schemes.

As previously described, applying a mixture model Peixoto (2021) to the vast ensemble of MCMC-generated partitions grouped structurally similar partitions into distinct clusters, revealing the dominant peaks within the solution space (see Section 4.3.2 for details). Each cluster represents a distinct *organizational scheme*, and the relative size of the cluster (*ω*) reflects that scheme’s prevalence, or probability of occurrence, within the landscape.

We observed four distinct organizational schemes within the solution landscape of the reference sample (Figure 7; see Section 4.1.6 for reference and bootstrap definitions). To determine whether these schemes are generalizable, we tracked them across all 500 resamples using their probabilistic assignment matrices (*π*; see Section 4.3.2 for details). A Friedman test performed on their tracked occurrence probabilities (*ω*) revealed no significant differences across the schemes (*χ*^2^(3) = 3.05*, p* = 0.55).

**Figure 7:**
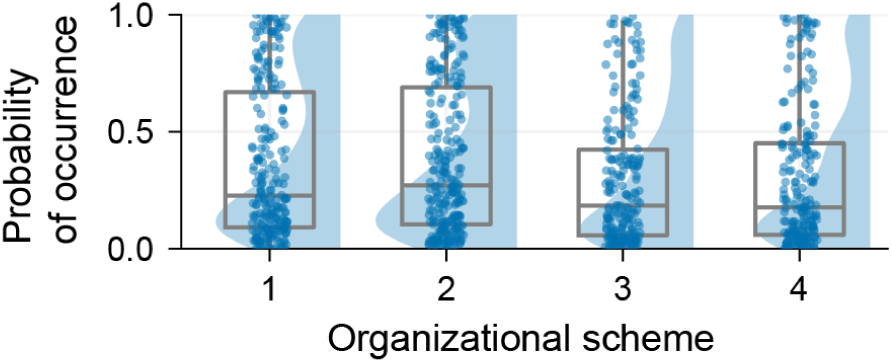
The resting-state functional connectome is characterized by multiple co-dominant organizational schemes. The probability of occurrence (*ω*) for each of the four distinct topological schemes tracked across the 500 bootstrap resamples. Despite representing different global topologies, there was no significant difference in the occurrence probabilities of these schemes (*χ*^2^(3) = 3.05, *p* = 0.55, Friedman test), demonstrating that the network’s underlying architecture is supported by an inherently multi-modal solution landscape rather than dominated by a single organizational scheme.

This lack of significant difference suggests that the resting-state functional architecture is not characterized by a single topological configuration. Instead, the functional connectome is inherently multi-modal, supported by a solution landscape of co-dominant organizational schemes that offer highly comparable explanations of the underlying network.

Notably, this inherent multi-modality was not observed when employing a traditional community detection approach. Evaluating the connectome using modularity maximization Newman and Girvan (2004); Newman (2006, 2016) revealed a unimodal landscape (see Supplement 5.3 for details). The modularity objective landscape is known to form a highly irregular plateau of structurally similar partitions Good et al. (2010). Applying our exact MCMC sampling and mixture model pipeline to this objective yielded only a single organizational scheme, suggesting that our mixture model framework effectively absorbs minor topological fluctuations into a unified cluster. Consequently, the multiple co-dominant schemes identified by our generative model reflect genuinely distinct network topologies rather than minor variations along a rugged plateau (which we investigate below).

We further verified that this inherent multi-modality is a robust property of the mouse functional connectome rather than an artifact driven by an outlier subject. A systematic leave-one-out analysis successfully recovered a multi-modal landscape across all nine leave-one-out iterations (*n* = 9). This result suggests that the presence of multiple co-dominant organizational schemes—a form of structural degeneracy—may represent a generalizable, population-level phenomenon (see Supplement 5.4 for details).

### 2.5 Structural variation within communities underlies the multiple organizational schemes

Having established that the functional connectome is characterized by multiple co-dominant organizational schemes, we next investigated how individual communities structurally manifest as distinct ROI assignment patterns across the posterior solution landscape to support these schemes.

As previously defined, a community *c* is the *c*^th^ column in the probabilistic assignment matrix (*π*) of each organizational scheme, representing the probabilistic assignments of the ROIs to that community. For each community, we aggregated the ROI assignment vectors (the corresponding columns from the *π* matrices) across all schemes and all bootstrap resamples. We then applied a mixture model (Gaussian Mixture Model Murphy (2012, 2022, 2023)) that grouped similar assignment vectors into distinct clusters (see Section 4.3.5 for details). Each cluster represents a distinct pattern of ROI assignments of the community, and the relative size of the cluster reflects that ROI assignment pattern’s prevalence (*ω*) in the solution landscape. To pinpoint which regions drive this structural variation, we evaluated the variance of each ROI’s assignment probabilities across the distinct patterns using non-parametric statistical testing (Mann-Whitney U or Kruskal-Wallis; see Section 4.3.5 for details), allowing us to label ROIs as either structurally stable or structurally variable.

Applying this clustering approach to each community revealed a spectrum of structural variation. Not all communities exhibited this variation; several possessed a single, stable ROI assignment pattern (e.g., the Cingulo-Prefrontal community, C.04, as shown in Figure 5). Furthermore, among the com munities that did exhibit structural variation, certain communities possessed hemispheric splits (e.g., the Medial-Temporal community, C.06). The remaining communities (C.01, C.02, C.03), however, exhibited coordinated bilateral variation in their assignment patterns, reflecting architectural trade-offs along distinct functional axes.

Figure 8 illustrates the distinct ROI assignment patterns of the Somatomotor community (C.02). We observed that this community primarily exhibited two co-dominant assignment patterns (top row). In the first pattern (left; *ω* = 53%), primary motor regions were assigned to the community with high probability (0.9) alongside somatosensory regions, yielding high structural resemblance to the canonical somatosensory system (R.01). In the second pattern (right; *ω* = 46%), however, the assignment probabilities of the primary motor regions were substantially reduced (0.5; *p* ≪ 0.05, Mann-Whitney U test, *α* = 0.05; see Section 4.3.5) with respect to the first pattern. This structural shift is isolated in the differential map (bottom row). This structural variation potentially reflects the underlying architectural signature of passive sensory monitoring without concurrent motor engagement.

**Figure 8:**
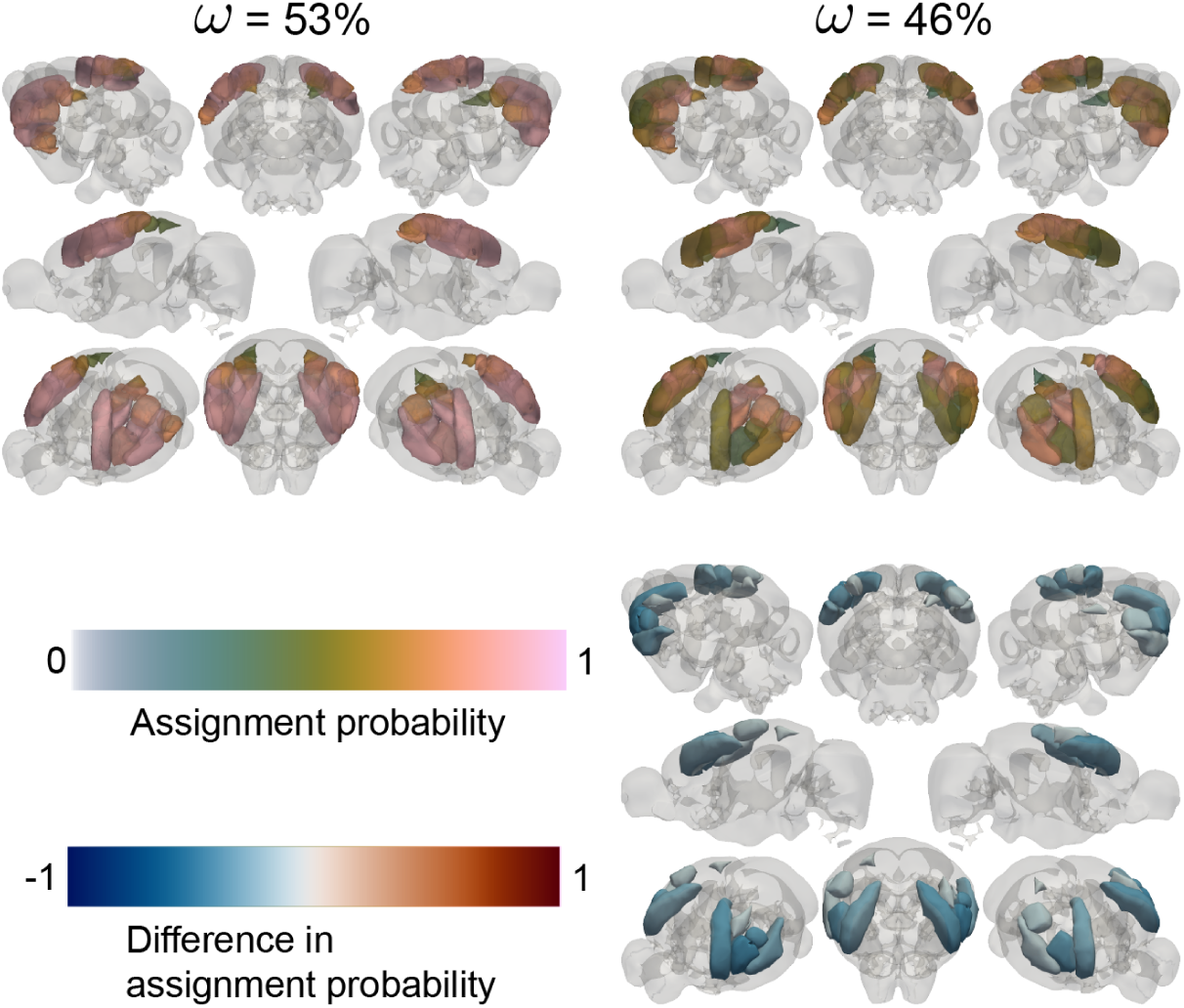
Structural variation within the Somatomotor community reveals variable motor region engagement. (Top row) The ROI assignment probabilities in the two patterns exhibited by the Somatomotor community (C.02), along with their prevalences (*ω*) in the solution landscape. The first pattern (left) integrates somatosensory and primary motor regions, while the second pattern (right) exhibits a more sensory architecture. (Bottom row) Differential map highlighting the directional change in assignment probabilities from the first pattern to the second pattern for structurally variable ROIs (*p <* 0.05). The pronounced negative difference (blue) mainly isolates primary motor areas from the structurally stable somatosensory regions.

Figure 9 illustrates the distinct ROI assignment patterns of the Olfactory-Prefrontal-Thalamic community (C.01). We observed that this community primarily exhibited three co-dominant assignment patterns (top row). In the most prevalent pattern (left; *ω* = 54%), the olfactory bulb was assigned to the community with high probability (0.77) alongside moderate assignments from the prefrontal cortex and thalamus, establishing a baseline sensory-associative axis. In the alternative patterns, however, the associative regions exhibit structural shifts. In the second pattern (middle; *ω* = 25%), prefrontal assignment probabilities substantially increase (0.74) while thalamic probabilities decrease (0.33) with respect to the first pattern. Conversely, in the third pattern (right; *ω* = 21%), prefrontal assignments diminish with respect to the first pattern. These structural shifts are isolated in the differential maps (bottom row; Kruskal-Wallis test, FDR-corrected, *α* = 0.05; see Section 4.3.5). This structural variation likely reflects the underlying architectural signature of variable involvement of higher-order regions during associative processing.

**Figure 9:**
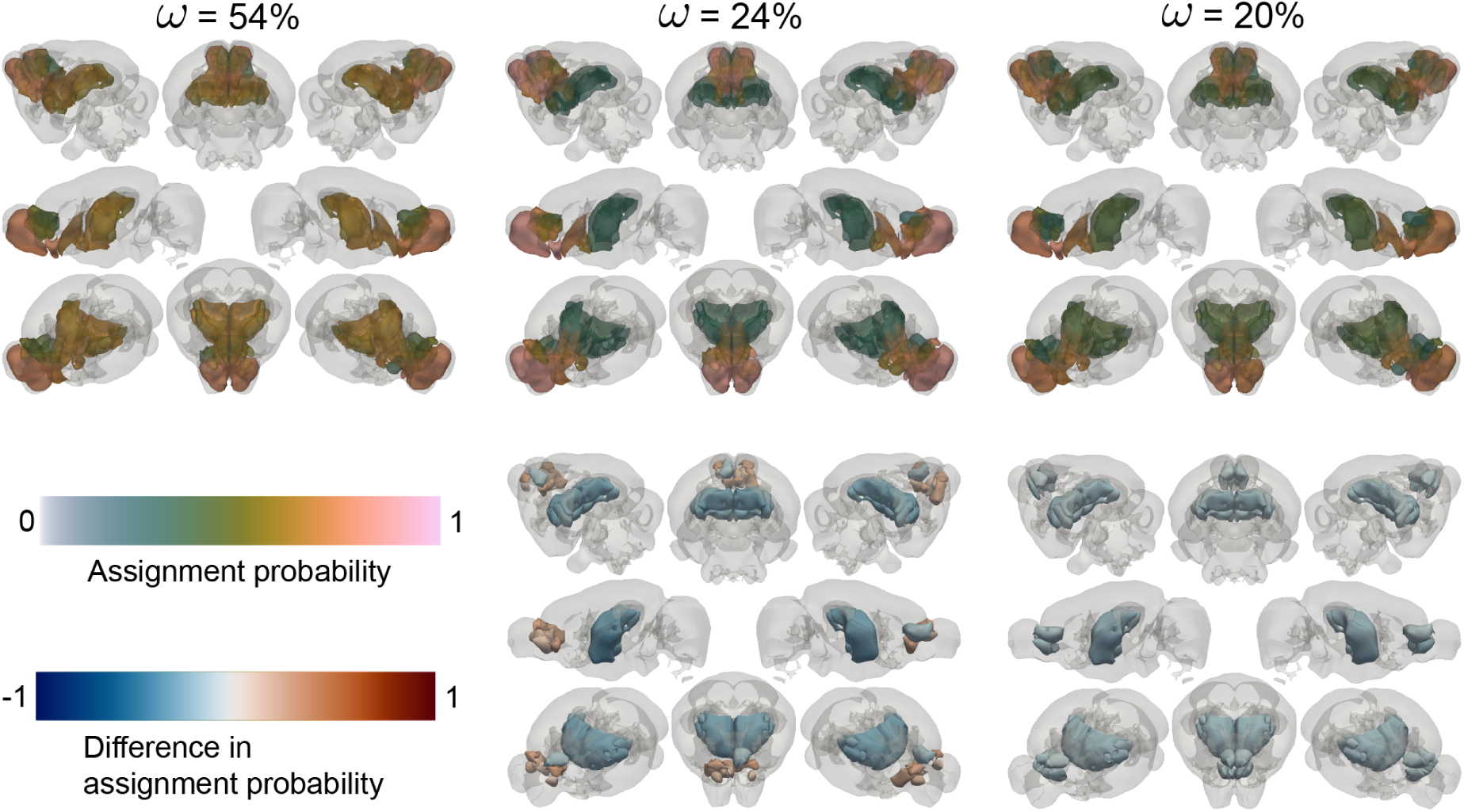
Structural variation within the Olfactory-Prefrontal-Thalamic community reveals variable engagement of higher-order regions. (Top row) The ROI assignment probabilities in the three patterns exhibited by the Olfactory-Prefrontal-Thalamic community (C.01), along with their prevalences (*ω*) in the solution landscape. The most prevalent pattern (left) establishes a baseline sensory-associative axis, while the alternate patterns exhibit either increased (middle) or decreased (right) prefrontal coupling. (Bottom row) Differential maps highlighting the directional change in assignment probabilities from the first pattern to the second (middle) and third (right) patterns for structurally variable ROIs (*p <* 0.05, FDR-corrected). The maps isolate divergent, bidirectional structural shifts of the prefrontal and thalamic regions relative to the first pattern.

Figure 10 illustrates the distinct ROI assignment patterns of the Fronto-Insular-Striatal community (C.03). We observed that this community primarily exhibited three distinct, co-dominant assignment patterns (top row). In the first pattern (left; *ω* = 36%), the dorsal striatum and agranular insula were assigned to the community with high probability (0.65), establishing a strong cortico-subcortical axis. In the alternative patterns, however, the community exhibits fundamentally different cortical affiliations. In the second pattern (middle; *ω* = 32%), assignment probabilities for secondary motor regions were substantially higher (0.58) with respect to the first pattern. Conversely, in the third pattern (right; *ω* = 31%), prefrontal assignment probabilities were substantially higher (0.47) while dorsal striatum assignments were lower (0.54) with respect to the first pattern. These structural shifts are isolated in the differential maps (bottom row; Kruskal-Wallis test, FDR-corrected, *α* = 0.05; see Section 4.3.5). The vast majority of ROIs in this community were structurally variable. This structural variation likely reflects the underlying architectural signature of a routing hub, capable of supporting alternative couplings between motor planning and executive control circuits to meet varied behavioral demands.

**Figure 10:**
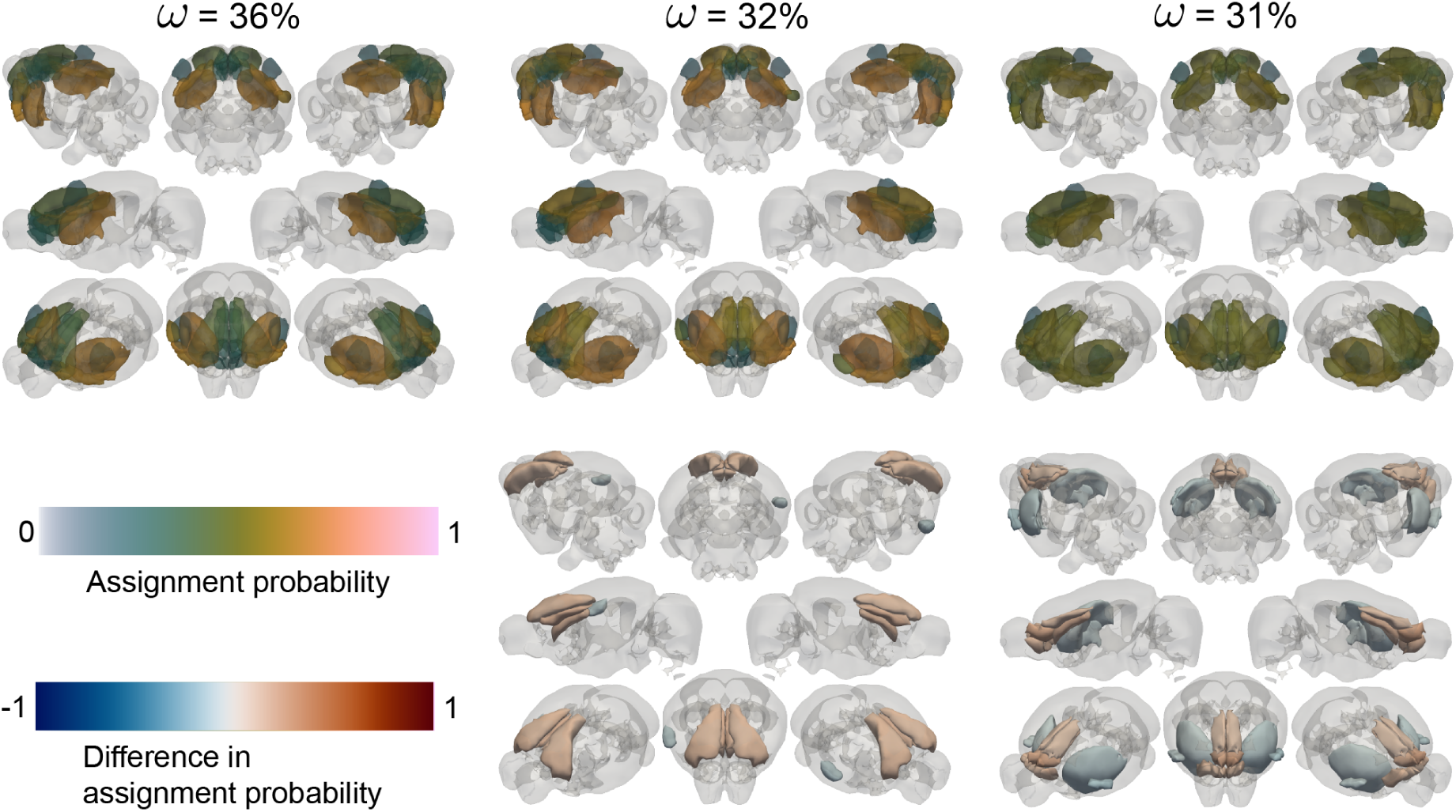
Structural variation within the Fronto-Insular-Striatal community reveals variable cortico-striatal architecture. (Top row) The ROI assignment probabilities in the three co-dominant patterns exhibited by the Fronto-Insular-Striatal community (C.03), along with their prevalences (*ω*) in the solution landscape. The first pattern (left) establishes a baseline striatal-insular axis, while the alternate patterns exhibit higher coupling with secondary motor (middle) or prefrontal (right) regions. (Bottom row) Differential maps highlighting the directional change in assignment probabilities from the first pattern to the second (middle) and third (right) patterns for structurally variable ROIs (*p <* 0.05, FDR-corrected). The maps isolate divergent structural shifts of motor and prefrontal cortices relative to the first pattern.

Together, these findings reveal that structural variation is not uniformly expressed across the connectome, but instead follows a systematic architectural pattern across communities. Primary and anatomically constrained communities exhibited structurally stable organization, whereas higher-order and integrative communities displayed substantial variation in their assignment patterns. This structural variation was organized along interpretable functional axes, such as sensorimotor engagement or prefrontal-thalamic coupling. These community-level variations result in the multi-modal solution landscape at the whole-brain level: distinct organizational schemes arise from coordinated structural shifts within variable communities, while structurally stable communities anchor the overall architecture. In this sense, the functional connectome appears to be composed of structurally stable cores and variable boundaries, enabling multiple co-dominant configurations to coexist.

## 3 Discussion

A central goal of systems neuroscience is to understand the principles that govern large-scale functional organization in the brain. In this study, we addressed two related challenges that have complicated this goal: evaluating the extent to which different organizational principles are compatible with functional connectivity data, and determining whether the resulting network organization is best characterized by a single partition or by a more heterogeneous solution landscape.

To address the first challenge, we employed a Bayesian generative modeling framework that makes explicit the assumptions underlying different organizational hypotheses. By comparing these models on a common information-theoretic scale (total description length), we found that within the set of models considered here, a non-degree-corrected hierarchical formulation provides the most parsimonious (thus most compatible) description of the observed functional connectivity patterns, and the purely assortative formulation the least (see Section 2.1). This does not falsify the importance of assortativity — which is undeniably a fundamental principle of brain networks Betzel et al. (2018b) — but it demonstrates that enforcing pure assortativity alone may give an incomplete description of the system. We specifically focused our analysis on the middle level of this hierarchy because its spatial scale is highly comparable to canonical resting-state networks (RSNs). When examined in relation to these standard RSNs, the inferred communities showed a systematic reorganization: primary sensory systems were largely preserved as cohesive units, whereas higher-order associative systems were distributed across multiple communities (see Section 2.3). These results suggest that canonical RSN definitions may aggregate distinct substructures that can be further resolved by generative modeling.

To address the second challenge, we examined the assumption that the space of network partitions is well summarized by a single consensus solution. By characterizing the posterior distribution over partitions under the hierarchical model, we identified multiple distinct modes (organizational schemes — topological arrangements of nodes (ROIs) into communities) in the solution landscape (see Section 2.4). Statistical testing did not reveal a significant difference in the prevalence of these modes across bootstrap resamples, indicating that no single organizational scheme dominates the landscape within the resolution of the present analysis. Further analysis showed that differences between modes arise from structured variability in the assignment of specific regions, with some communities exhibiting stable composition and others showing systematic variation across modes (see Section 2.5).

Taken together, these results suggest that, within the present modeling framework, the functional connectome is not adequately described by a single partition. Instead, the data support a set of alternative partitions that differ in the assignment of specific regions while preserving broader organizational structure. This perspective provides a potential explanation for variability across previous studies, which may arise when a heterogeneous solution landscape is forced into a single consensus partition.

### 3.1 Inferred communities refine canonical networks

The communities inferred under the hierarchical model exhibit clear spatial organization, forming contiguous and bilaterally symmetric patterns despite the absence of explicit spatial priors. These features indicate that the generative model captures structured aspects of the functional connectivity data. However, when evaluated against a canonical resting-state network (RSN) atlas Zerbi et al. (2015), the inferred communities reveal systematic differences in how functional boundaries are drawn, particularly between sensory and associative cortices.

Bidirectional geometric analysis demonstrated that primary sensory systems remain highly cohesive. The canonical somatosensory network, for instance, mapped closely onto the inferred Somatomotor community, indicating a strong correspondence between this RSN and the community identified by the model. In contrast, higher-order association networks were distributed across multiple inferred communities. For example, the canonical limbic system was partitioned across the Olfactory-Prefrontal, Medial-Temporal, and Striato-Amygdalar communities, suggesting that this RSN encompasses multiple distinct substructures under the present framework.

Rather than invalidating canonical networks, these differences highlight a distinction in resolution and methodology. The RSN atlas was derived from post-hoc clustering of independent components Zerbi et al. (2015), providing a highly useful and broad heuristic for functional macro-regions. Conversely, our generative framework infers community structure directly from the data under an explicit probabilistic model. As a result, the generative model refines these broad territories, suggesting that some canonical RSNs, especially those associated with associative processing, may actually reflect aggregate structures composed of multiple sub-components. This observation is consistent with perspectives that emphasize distributed organization in large-scale brain networks Pessoa (2022, 2023), though further work is required to determine how these organizational differences relate to underlying neural or functional processes.

### 3.2 Multiplicity reveals structured variability across communities

Mapping the posterior distribution of the optimal hierarchical model revealed that the functional connectome is characterized by multiple distinct modes in the space of network partitions (the solution landscape). In contrast, evaluating the connectome using modularity maximization yielded a largely uni-modal landscape. Again, this does not falsify the importance of assortativity, but it demonstrates that enforcing a pure assortativity as the sole organizational rule provides an incomplete description that obscures underlying multiplicity. More flexible generative models admit a broader range of topological configurations that are equally consistent with the observed data.

This contrast highlights an important theoretical nuance regarding Bayesian model selection Gel-man et al. (2013); Murphy (2023). From an information-theoretic perspective, the total description length (TDL) inherently penalizes the entropy of the posterior distribution, favoring models with unimodal solution landscapes. Ideally, if a generative model perfectly captured the true underlying process of the data, it would yield a unique and unambiguous partition. The fact that our optimal model yields a multi-modal landscape indicates a degree of structural uncertainty, reflecting a mild “mis-fit” between the generative constraints of hierarchies and the profound complexity of the functional connectome Peixoto (2021); Grunwald (2004). However, as demonstrated by our analysis, possessing a unimodal landscape is not a guarantee of a more compatible model. Modularity maximization forced an almost entirely unimodal landscape, yet it provided the least compatible description of the data. Therefore, while this multiplicity serves as a strong motivation to explore even more advanced generative models in the future, it currently provides the most accurate and honest reflection of the connectome’s architecture (under our modeling choices).

To quantify how there modes differ, we analyzed the variability or regional assignments across the identified modes. This analysis revealed that variability is not uniformly distributed across the connectome, but instead follows a structured anatomical pattern. Some communities exhibited highly consistent assignments across modes, indicating stable organization. In contrast, other communities displayed substantial variability in their regional composition. For example, the Somatomotor community exhibited relatively constrained variability, while communities such as the Fronto-Insular-Striatal and Olfactory-Prefrontal-Thalamic systems showed broader variability, with systematic changes in cortical and subcortical affiliations. These patterns suggest that differences between modes arise from coordinated reassignments of subsets of regions, rather than from random fluctuations or global reorganization.

Importantly, these results must be interpreted within the scope of the statistical framework employed here. The identified modes reflect distinct high-probability partitions under the hierarchical generative model applied to a static functional connectivity network. As such, they characterize alternative ways of clustering the same time-averaged data from anesthetized animals, rather than directly corresponding to distinct functional states or dynamic information routing. Future work utilizing time-varying functional connectivity Lurie et al. (2020), or temporal independent component analysis Xu et al. (2022) will be required to determine whether these alternative static partitions relate to transient temporal fluctuations.

### 3.3 Methodological considerations

While our Bayesian framework identifies a non-degree-corrected hierarchical architecture as the most compatible organizational principle, we do not claim this represents the absolute ground truth of the brain. In generative modeling, model selection is inherently relative to the specific hypothesis space evaluated Murphy (2022, 2023). It remains possible, and is encouraged, that future theoretical advances will propose even more sophisticated generative principles — ones that do not exist yet — that surpass discrete hierarchies, potentially including time-varying generative rules capable of modeling fMRI time series data directly (for example see Peixoto (2024)).

A secondary consideration involves our specific model preferences. For instance, while the non-degree-corrected models were preferred over degree-corrected variants — a result that could intuitively suggest “rich-club” topographies where highly connected hubs group together — this preference must be interpreted with caution. Our network construction retained the top 20% of edges, a thresholding procedure that imposes strict constraints on the degree distribution. Furthermore, BOLD-derived correlation matrices possess inherent degree homogeneity Zalesky et al. (2012); Van Den Heuvel et al. (2017); Fornito et al. (2016). Without extensive sensitivity analyses, we cannot definitively conclude that this model preference reflects true neurobiological rich-club organization rather than an artifact of these methodological constraints.

Finally, three additional methodological caveats contextualize our findings. First, functional MRI relies on the BOLD signal, which introduces spatial and temporal blurring that can alter the topology of functional boundaries Vafaii et al. (2024). Consequently, the exact spatial configurations of our inferred communities are inevitably influenced by vascular architecture.

Second, the data were acquired from anesthetized mice. General anesthesia, and its specific dosage, globally depresses spontaneous cortical activity Bukhari et al. (2017); Barttfeld et al. (2015). Therefore, the structural multiplicity reported here strictly characterizes the anesthetized connectome. Future investigations in awake, behaving animals — and utilizing direct mesoscopic neural readouts like wide-field calcium imaging Lake et al. (2020) — are required to fully characterize the topological multiplicity of the brain.

Third, our results inherently depend on the choice of spatial parcellation. While our generative models received no spatial priors during community inference, the definition of the network nodes themselves relied on the Allen CCFv3 atlas. The size, shape, and anatomically predefined boundaries of these regions impose a foundational structural prior on the network topology. Future work should investigate whether deriving functionally-driven parcels directly from the time-series data (rather than relying on an anatomical reference atlas) alters the multiplicity of the solution landscape.

### 3.4 Conclusion

To summarize, our Bayesian generative modeling framework provides a principled re-evaluation of macroscopic functional organization in the mouse brain. By explicitly comparing competing topological hypotheses, we demonstrated that the functional connectome is optimally described by a hierarchical architecture, wherein broad associative networks are revealed to be composite structures composed of distinct sub-circuits. Furthermore, we demonstrated that the static functional connectome does not easily admit a singular consensus partition. Instead, the data support a multi-modal landscape of co-dominant organizational schemes, driven by structured variability of specific regional affiliations around structurally stable sensory anchors. Ultimately, these findings underscore a critical methodological shift for systems neuroscience: to accurately map the functional connectome, analytical frameworks must move beyond enforcing singular, rigid solutions and embrace the brain’s inherent structural variation.

## Acknowledgments

We thank Drs. Joel Greenwood, Omer Mano, and Paul Shamble from the Neurotechnology Core of the Yale Kavli Institute for Neuroscience for their technological expertise, and Dr. Xinxin Ge for assistance in acquiring the multimodal imaging data. We are grateful to Dr. Peter Herman for performing the head-plate implant surgeries and to all members of the Yale Multiscale Imaging and Spontaneous Activity in Cortex (MISAC) group for their contributions to conceptualizing and building the in-scanner WF-Ca^2+^ imaging apparatus. We owe a significant debt of gratitude to Dr. Tiago Peixoto for his invaluable guidance on the graph-tool library; his insights and generous support on the project’s discourse forum were instrumental in implementing the custom functions required for this work. We also thank Dr. Joyneel Misra, Dr. Xiaoyu Zhou, and Songtao Song for their helpful and insightful discussions throughout the development of this manuscript. This work was supported by National Institutes of Health grants R01 MH111424 and U01 NS120358, as well as the University of Maryland Brain and Behavior Initiative (BBI) Seed Grant (2022–2024).

## 4 Methods

### 4.1 Data acquisition, preprocessing, and structure

All procedures were performed following the Yale Institutional Animal Care and Use Committee (IACUC) and in agreement with the National Institute of Health Guide for the Care and Use of Laboratory Animals.

#### 4.1.1 Group and dataset overview

Mice were housed on a 12-hour light/dark cycle. Food and water were available ad libitum. Animals were 6-8 weeks old, 25-30g, at the time of the first imaging session. All groups were of mixed sex and shared a C57BL/6J background. Male CRE mice were selected from the offspring of parents with different genotypes; this avoided leaking of CRE expression. Briefly, the Ai162 genotype resulted from tTA and TITL-GCaMP6s (TIGRE1.0). Differences between TIGRE1.0 and 2.0 have been reported (Daigle et al., 2018) and TIGRE1.0 mice have been previously described (Lake et al., 2020).

#### 4.1.2 Surgical procedure for permanent optical access to the cortical surface and immobilization

All mice underwent a minimally invasive surgical procedure were an in-house built head-plate (acrylonitrile butadiene styrene plastic, TAZ-5 printer, 0.35mm nozzle, Lulzbot) was attached to the thinned, but intact, skull for permanent optical access to the cortical surface, necessary for WF-Ca2+ imaging, and immobilization. The surgical procedure has been described by us previously (Lake et al., 2020; Vafaii et al., 2024; O’Connor et al., 2022). Briefly, mice were initially anesthetized with isoflurane (5%, 70/30 medical air/O_2_) and head-fixed in a stereotaxic frame (KOPF, USA). Isoflurane was then reduced to 2%. An eye ointment was applied; meloxicam (2mg/(kg)body weight) administered subcutaneously and bupivacaine (0.1%) injected locally, at the incision site. After shaving the hair, the scalp was washed with betadine followed by ethanol 70%, (×3). The scalp was surgically removed, and the skull cleaned and dried. Antibiotic powder (Neo-Predef) was applied to the incision site, and isoflurane reduced (1.5%). Skull-thinning of the frontal and parietal skull plates was performed with a hand-held drill (FST), diameter of the tips 1.4mm and 0.7mm. Superglue (Locite) was applied to the exposed skull, followed by transparent dental cement C&B Metabond (Parkell). The head-plate was attached to the dental cement before solidification. The head-plate had a double-dovetail plastic frame with a microscope slide hand-cut to match the size and shape of the mouse skull (Luo and Constable, 2022).

#### 4.1.3 Imaging protocol

These data belonged to a dual-imaging dataset described elsewhere (Mandino et al., 2025a).

Mice were allowed 1-week to recover post-surgery before undergoing three simultaneous WF-Ca2+ and BOLD-fMRI sessions on an 11.7T MRI scanner (Bruker, Billerica, MA) using custom MRI-compatible WF-Ca^2+^ imaging equipment (Lake et al., 2020). The interval between each imaging sessions was at least 1-week.

Data were acquired as described previously (Lake et al., 2020; Vafaii et al., 2024) using ParaVision 6.0.1. Mice were scanned whilst lightly anesthetized with isoflurane and free breathing (0.5-0.75%, 70/30 medical air/O2). Body temperature was continuously monitored (Spike2, Cambridge Electronic Design Limited), and maintained with a circulating water bath. Imaging data and physiological recordings were synchronized (Master-8 A.M.P.I., Spike2 Cambridge Electronic Design Limited). A gradient-echo, echo-planar-imaging (GE-EPI) sequence with a 1.0s repetition time (TR) and 9.1ms echo time (TE) was used with an isotropic resolution of 0.4×0.4×0.4mm3. Twenty-eight slices yielded near whole-brain coverage. Each functional run was comprised of 600 volumes or 10mins of data. On average, 7 functional runs were acquired per mouse per session. Three included a unilateral LED-stimulation whilst 4 were acquired in the resting-state (no external stimuli were presented). Here, we consider the resting-state data only.

In addition to the functional data, 4 structural images were acquired during each imaging session for data registration purposes (Lake et al., 2020; Vafaii et al., 2024). (1) A multi-spin-multi-echo (MSME) image sharing the same FOV as the functional data with a TR/TE of 2500/20ms, 28 slices, two averages, and resolution of 0.1×0.1×0.4mm^3^ in 10mins and 40s. (2) A whole-brain isotropic MSME image with a TR/TE of 5500/20ms, 78 slices, two averages, and resolution of 0.2×0.2×0.2mm^3^ in 11mins and 44s. (3) A fast-low-angle-shot (FLASH) time-of-flight (TOF) angiogram, with a TR/TE of 130/4ms, resolution of 0.05×0.05×0.05mm^3^ and FOV of 2.0×1.0×2.5cm^3^ in 18mins. (4) A FLASH image of the angiogram FOV, with a TR/TE of 61/7ms, four averages, and resolution of 0.1×0.1×0.1mm^3^ in 11mins and 24s. Each average of images (1) and (2) were interleaved with functional scans such tha the functional data were acquired at approximately evenly spaced intervals throughout each imaging session.

#### 4.1.4 MRI data registration and preprocessing

BOLD-fMRI data processing was performed using RABIES (Rodent automated BOLD improvement of EPI sequences (Desrosiers-Gŕegoire et al., 2024)) v0.4.8 (https://rabies.readthedocs.io/en/stable/). All resting-state functional and isotropic MSME structural scans (from all imaging sessions) were input and processed together. Within native subject space, the BOLD-fMRI timeseries were slice time corrected, and head motion was estimated. Structural images were corrected for inhomogeneities (N3 nonparametric nonuniform intensity normalization). The within-sample anatomical template, created by averaging all structural images following non-linear registration, was registered to our in-house template (using non-linear registration), which has been previously registered to the Allen Atlas reference space CCfv3 (Vafaii et al., 2024; Wang et al., 2020; Mandino et al., 2025b).

For each BOLD-fMRI scan, a representative mean image (averaged across time) was derived. These data were corrected for intensity inhomogeneities and non-linearly registered to the corresponding isotropic structural MSME image of each mouse acquired during each imaging session. This minimized distortions caused by susceptibility artifacts. Then, the BOLD-fMRI data were moved to common space using four concatenated transformations: (1) framewise rigid head motion correction, (2) representative mean BOLD-fMRI to individual mouse/session isotropic MSME image, (3) individual mouse/session isotropic MSME image to within-dataset template, and (4) within-dataset template to out-of-sample in-house template. As part of this transformation, the data were resampled to the template resolution (0.2×0.2×0.2mm^3^). After this step, the Allen Atlas and BOLD-fMRI data resided in the same space. Registration performance was visually inspected using the RABIES quality control report. Following timeseries normalization to common space, the six-parameter motion estimates were regressed from the timeseries and used to compute framewise-displacement. Frame scrubbing for motion using a conservative 0.075mm threshold was applied. Data were filtered 0.008-0.2Hz and 30 timepoints discarded from the beginning and end of each run to avoid filter-related edge artifacts. Cerebrospinal fluid and white matter signals were regressed, and the data were smoothed with 0.4mm sigma.

#### 4.1.5 Data structure and network construction

The dataset possesses a hierarchical structure: data were collected from 9 mice, each undergoing three longitudinal imaging sessions consisting of four 10-minute resting-state fMRI runs (108 total runs). The input to our generative modeling pipeline is a single, representative group-level functional network derived from this data.

First, we computed Pearson correlation matrices for each fMRI run. To account for variations in run length due to motion scrubbing, we aggregated these run-level matrices into subject-level matrices, and subsequently into a single group-level matrix, using a timepoint-weighted averaging procedure (see 5.1.1 for normalization and Fisher *r*-to-*z* transformation details) (Zar, 1999; Power et al., 2014). To create a sparse, undirected graph suitable for network analysis, we applied a proportional threshold to this group-level matrix, retaining the top 20% of correlation values. This thresholding exclusively isolates the strongest positive functional relationships, preserving the core topological structure of the brain while eliminating spurious, low-magnitude noise. The resulting matrix was binarized to create the final functional network used for model fitting.

#### 4.1.6 Bootstrap resampling

To ensure our subsequent topological findings were robust to population variance and representative of the true underlying architecture, we embedded this network construction procedure within a bootstrap resampling framework. Specifically, we sampled the 9 subjects with replacement 500 independent times. For each of these 500 bootstrap resamples, we executed the weighted averaging and thresholding procedure described above to generate a unique bootstrap-resample-level functional graph.

Consequently, our dataset for all subsequent Bayesian inference consisted of an ensemble of 500 independent bootstrap graphs. The entire analytical pipeline was executed independently on each of these graphs, allowing us to generate population-level confidence intervals for all downstream analyses.

For convenience, we refer to the original sample of mice as the *reference sample* to differentiate it from all bootstrap resamples.

### 4.2 Framework of Statistical inference

#### 4.2.1 Bayesian generative modeling

To infer the community arrangements underlying the observed functional networks, we adopted a Bayesian generative modeling approach. This statistical framework (Peel et al., 2022) allows us to formally test competing hypotheses about the principles of brain network organization, moving beyond heuristic pattern description to rigorous statistical inference. The core idea is to reframe the central question from “what communities exist?” to “how likely is it that a specific underlying community structure *generated* our observed network?”

This framework relies on a generative model, which acts as a probabilistic recipe for how a network is constructed based on a latent community structure and a set of organizational rules. Using Bayesian inference, we invert this process to calculate the *posterior* probability of any proposed community partition. This calculation balances the *prior* (the assumptions of a given organizational hypothesis, such as assortativity) against the *likelihood* (the probability of observing the actual brain network if that hypothesized structure were true).

Crucially, to identify the optimal community structure without arbitrarily fixing the number of communities in advance, we framed this probabilistic inference as an information-theoretic optimization problem. The posterior probability of a partition can be rewritten in terms of an information theoretic quantity called *description length* Σ, that quantifies the amount of bits required to describe the network in terms of the partition. Following the Minimum Description Length (MDL) principle (Rissanen, 1998; Grunwald, 2004), the most probable explanation for a dataset is the one that provides the most efficient compression of that data. Therefore, our inference algorithm searches for the community partition that minimizes the description length (Σ) of the network. A shorter description length indicates a more parsimonious, and computationally more probable, model of the network’s true organization (see 5.1.2 for the complete mathematical formulation).

#### 4.2.2 Generative models of brain organization

We instantiated our Bayesian framework using several variants of the stochastic block model (SBM), specifically as formulated in (Peixoto, 2019). At its core, an SBM is a generative model for networks which posits that nodes are organized into groups, or “blocks”, and that the probability of a connection between any two nodes depends solely on the blocks to which they belong. All SBM variants and subsequent model fitting procedures were implemented using the graph-tool Python library (Peixoto, 2014b).

Each SBM variant represents a concrete, testable hypothesis about the organizational principles shaping brain networks. We evaluated three primary structural architectures (see 5.1.3 for complete mathematical formulations):

- **Standard SBM (s-SBM)** (Peixoto, 2017): This model serves as a flexible foundation. It partitions nodes into groups but imposes no prior constraints on how these blocks connect. It can capture a wide range of topographies, from segregated assortative modules to integrative core-periphery or bipartite arrangements (Betzel et al., 2018a).
- **Hierarchical SBM (h-SBM)** (Peixoto, 2014c): Brain networks are thought to be organized across multiple scales. The h-SBM explicitly models this by assuming communities are nested within larger, higher-order communities, recursively partitioning the network to reveal a multi-scale hierarchy.
- **Overlapping SBM (o-SBM)** (Peixoto, 2015): To account for brain regions participating in multiple functional systems, the o-SBM partitions the connections (half-edges) rather than the nodes themselves. This naturally allows a node’s “membership” to be split as a mixture across several functional groups.

Furthermore, for each of these three architectures, we tested both non-degree-corrected (nd-) and degree-corrected (dc-) variants. The degree-corrected formulation is a cornerstone of modern network science (Karrer and Newman, 2011). Non-degree-corrected models assume all nodes within a community have a statistically similar number of connections. In contrast, degree-corrected models decouple a node’s overall connection strength from its community assignment.

This distinction pits two competing neurobiological hypotheses against each other. A degree-corrected model treats a region’s “hubness” (Bullmore and Sporns, 2009) as an intrinsic local property separate from its community. Conversely, a non-degree-corrected model assumes connectivity is a feature of the community itself (e.g., all nodes in a “hub community” form a highly connected “rich-club”). Comparing these variants reveals which organizational principle better explains the empirical functional architecture.

#### 4.2.3 The modularity benchmark and its generative counterpart

A widely used method for brain community detection is modularity maximization (Newman, 2006; Sporns and Betzel, 2016), which identifies partitions with dense within-community connections and sparse between-community connections. While powerful, this is a descriptive heuristic that optimizes a single score (modularity), rather than a generative model explaining network formation.

However, modularity maximization is mathematically equivalent to maximum likelihood estimation of a special case of the degree-corrected SBM, known as the degree-corrected Planted Partition model (which we refer to as the **assortative SBM (a-SBM)**; see 5.1.4) (Newman, 2016; Zhang and Peixoto, 2020).

This equivalence implies both methods seek to explain the exact same underlying principle: strict assortativity. Yet, they do so fundamentally differently. Modularity maximization will forcefully return a high-scoring assortative solution regardless of whether that configuration is genuinely supported by the data. In contrast, the a-SBM assigns probabilities to configurations and penalizes models that lack statistical evidence. This allows us to directly test the validity of the assortative assumption: if modularity maximization yields stable solutions, but the a-SBM fails to identify a coherent configuration, it indicates that modularity is overfitting to incidental noise rather than detecting true structural rules.

#### 4.2.4 Model fitting: MCMC simulations

The posterior distribution over all possible network partitions—which we refer to as the “solution landscape”—is incredibly vast and high-dimensional, making it computationally impossible to solve analytically. Therefore, we approximated this posterior distribution using Markov chain Monte Carlo (MCMC) sampling (Gelman et al., 2013).

At its core, MCMC is an algorithm that starts at a random partition and explores the landscape by iteratively proposing changes to the community assignments, preferentially accepting changes that improve the model’s fit. We utilized a highly advanced MCMC algorithm implemented in the graph-tool Python library (Peixoto, 2014b) specifically curated for graph structures. This algorithm explores the landscape using two smart, topology-driven mechanisms: first, it uses *neighborhood-guided* proposals, as opposed to random proposals, relocating a node to a community where it already has existing connections (Peixoto, 2014a). Second, to escape getting trapped in local minima, it utilizes macroscopic *merge-split* proposals, periodically merging two entire communities together or splitting one in half to make massive leaps across the solution landscape (Peixoto, 2020).

This guided “random-walk” yields an ensemble of highly probable partitions that accurately represent the true posterior distribution (see 5.1.5 for exact acceptance criteria). For each generative model evaluated, we ran five independent MCMC chains, each for 100,000 steps. Running five chains is a standard, robust practice in Bayesian inference (Gelman et al., 2013); it provides a mathematically sufficient number of pairwise comparisons to ensure the sampler has fully explored the global landscape from different starting points, without incurring prohibitive computational costs.

#### 4.2.5 Validating model faithfulness: MCMC convergence diagnostics

Before comparing different models (that is, organizational principles), it is crucial to ensure that each model provides a “faithful” explanation of the data. This diagnostic step is grounded in a core principle of Bayesian inference: a mathematically well-specified model whose assumptions are compatible with the network’s underlying structure should produce a single, coherent posterior landscape (Peixoto, 2023). While this landscape may be complex, it should be continuously connected and thoroughly explorable.

The convergence of multiple, independent MCMC chains onto this exact same landscape serves as a direct test of the model’s coherence. If independent chains initialized from different starting points consistently find and explore the same high-probability regions, it indicates that the model’s assumptions align with the data. Conversely, if chains become permanently trapped in irreconcilable, disconnected regions, it strongly indicates that the model’s structural assumptions are incompatible with the data, causing the posterior landscape to pathologically fracture.

To assess this, we monitored the description length (Σ) distributions sampled by each independent chain. Convergence was evaluated both qualitatively (via distribution overlap) and quantitatively using the Kolmogorov-Smirnov (KS) distance (see 5.1.6).

As chains converge onto a unified landscape, their sampled Σ distributions should become nearly identical. To formalize this, we utilized the KS distance as a strict geometric tolerance limit. We required that the absolute maximum difference between the distributions of any two chains drop below a standard threshold of 0.2 (Gibbons and Chakraborti, 2011) (see 5.1.6 for details). Models that failed to achieve this strict spatial convergence across all independent chains were considered unfaithful representations of the data and were excluded from all subsequent topological analyses.

### 4.3 Analysis Procedures

#### 4.3.1 Identifying the organizing principle: Model selection via Total Description Length

After identifying the set of faithful models that successfully achieved MCMC convergence, we performed a formal model comparison to determine which organizational principle provided the most parsimonious explanation of the mouse functional connectome. We evaluated the competing models based on their model evidence—the probability of observing the network data integrated over the entire ensemble of possible partitions generated by the model.

For practical computation and information-theoretic interpretation, we calculated the negative logarithm of this evidence, termed the Total Description Length (TDL). Following the Minimum Description Length (MDL) principle (Rissanen, 1998), TDL quantifies the total number of bits required to compactly represent the network data using the specific structural rules of the model. The model yielding the lowest TDL provides the most efficient data compression and is formally considered the optimal, most probable generative explanation.

Intuitively, TDL acts as a rigorous mathematical formulation of Occam’s razor (see 5.1.7 for exact derivation). TDL integrates over the entire solution landscape (that is, the posterior distribution of partitions), rather than evaluating a model solely on a single “best-fit” partition. Crucially, while TDL penalizes unnecessary model complexity, it does not artificially punish a model for producing a multi-modal solution landscape. It calculates a precise balance between the model’s average goodness-of-fit and the entropy (the statistical uncertainty, or “spread”) of its solution landscape. If the data inherently contains multiple valid organizational schemes, TDL accommodates this multiplicity, effectively preferring a model that accurately captures the degenerate solution landscape over a simpler model that underfits the data by forcing a single, unimodal consensus Peixoto (2021).

For each network, TDL was calculated for each independent MCMC chain using the graph-tool Python library (Peixoto, 2014b) and averaged across chains to provide a single, stable estimate. By comparing these final TDL values, we objectively adjudicated between our competing structural hypotheses, testing whether the resting-state functional architecture is best governed by simple assortative modules, overlapping communities, or a nested hierarchy, thereby selecting the single optimal generative framework for all subsequent topological analyses.

#### 4.3.2 Characterizing global multiplicity: organizational schemes of the whole brain

##### Partition alignment

A critical step before analyzing the partitions generated by MCMC sampling is alignment. Because community labels are arbitrarily assigned during the sampling process, equivalent functional groups might receive different integer labels across different samples—a well-known statistical issue termed “label switching” (Stephens, 2000). To resolve this, we utilized the random label model (Peixoto, 2021). Conceptually, this model posits that all sampled partitions are variations of a single, hidden “canonical” labeling template, but with their integer labels randomly permuted. By treating the observed assignments as random permutations, the algorithm mathematically “un-shuffles” the labels of each individual sample. It maximizes the node-for-node overlap between partitions, explicitly matching and re-numbering groups of nodes that consistently cluster together (see 5.1.8 for algorithmic details). This ensures that a given community label represents the exact same community across the entire ensemble.

##### Multimodal mixture model

We hypothesized that the posterior distribution of the functional connectome might *not* form a single, narrow peak, but rather a complex, multi-modal landscape representing multiple distinct topological configurations. To explicitly test this hypothesis and discover potential multiplicity, we clustered the vast ensemble of aligned MCMC samples into a discrete, interpretable set of *K* distinct solution *modes* (Peixoto, 2021). Conceptually, this algorithm groups structurally similar network partitions together to identify the dominant “peaks” (or centers of mass) within the highly complex solution landscape.

The mixture model yields two key outputs for each mode *k*:

1. The **mode probability (***ω_k_*): The relative mass, or fraction of the overall posterior landscape, occupied by mode *k*, representing its statistical likelihood (that is, *_k_ ω_k_*= 1).
2. The **node-membership probability matrix (***π*^(^*^k^*^)^): A matrix defining the precise topological structure of the mode, where entries denote the probability of a specific ROI (a row of the matrix) belonging to a specific community (a column of the matrix).

This two-tiered summary quantifies both the structural uncertainty *within* a given organizational mode (via *π*^(*k*)^) and the statistical uncertainty *between* competing modes (via *ω_k_*).

##### Topological merging of redundant modes

The mixture model identifies statistically distinct peaks in the posterior landscape. However, two closely situated peaks might represent the exact same macroscopic topological state, differing only by minor noise or probability scaling localized to a few ROIs. To bridge the gap between strict statistical non-equivalence and true neurobiological redundancy, we performed a post-hoc structural merge of the identified modes.

Procedurally, we compared the overall topological “shape” of every mode against every other mode using a geometric similarity metric (cosine similarity). To guarantee we only merged modes that were genuinely redundant (and not simply similar by chance) we generated a random baseline by repeatedly shuffling the ROI labels. Modes that were significantly more similar to each other than to this spatially shuffled baseline were mathematically consolidated (see 5.1.8 for exact permutation and FDR-correction details). The resulting merged clusters, with their probabilities (*ω_k_*) summed together, represent the final, parsimonious set of macroscopic *organizational schemes* — the distinct ways in which the whole brain can functionally organize itself under a given organizational principle (e.g., hierarchical).

##### Aligning schemes across bootstrap resamples

Finally, this entire pipeline was executed independently on all 500 bootstrap resampled graphs. To enable group-level statistical testing, the organizational schemes identified in each bootstrap resample had to be mapped back to the reference schemes derived from the original, unresampled graph.

Procedurally, this involved taking the set of schemes found in a specific bootstrap resample and matching them one-to-one with the most structurally similar reference schemes. We achieved this tracking using the *linear-sum-assignment* Kuhn (1955); Munkres (1957); Jünger et al. (2010) algorithm that globally optimizes these pairings based on their topological overlap, ensuring we were reliably comparing structurally equivalent schemes across different resamples. As before, we verified that these algorithmic matches were statistically genuine by comparing them against spatially shuffled null models (see 5.1.8).

Ultimately, to determine whether the functional connectome exhibited a statistical preference for any specific organizational scheme, we extracted the tracked prevalence (*ω*) for each matched scheme across the 500 resamples. Because the prevalence values extracted from a single bootstrap resample represent a set of dependent probabilities that are mathematically constrained, we performed a non-parametric Friedman test to evaluate the null hypothesis that there is no significant difference in their occurrence probabilities across the schemes.

#### 4.3.3 Defining the reliable shape of communities

Before analyzing the fine-grained structural multiplicity of individual communities, we first rigorously define their topological shapes—that is, the set of ROIs that reliably constitute a community.

To establish a baseline structure for each bootstrap resample, we calculated an expected node-membership matrix. We computed the weighted average (proportional to *ω_k_*) of the node-membership matrices (*π*^(^*^k^*^)^) across all schemes identified within that resample. This step provides a single expected node-membership matrix that accounts for the structural uncertainty across schemes (see 5.1.9 for exact formulations).

With an expected node-membership matrix generated for each resample, our next step was to separate true member ROIs from high-variance background noise. We compiled the distribution of each ROI’s expected membership for a community across the 500 bootstrap resamples and tested it against a null hypothesis of zero affiliation. Only ROIs that exhibited a statistically significant (that is, non-zero) membership across the population—after rigorous False Discovery Rate (FDR) correction (see 5.1.9)—“survived” this thresholding process.

ROIs that failed to survive this test were classified as incidental noise for that specific community and masked out. The surviving ROIs formally defined the reliable spatial boundaries of the community.

#### 4.3.4 Biological contextualization of inferred communities: Resemblance to canonical resting-state networks

Having established the strict spatial boundaries of our inferred communities, our next objective was to biologically contextualize these communities. We achieved this by evaluating their spatial alignment with a canonical set of functional networks.

##### Canonical functional systems

We adopted the resting-state network (RSN) atlas defined by Zerbi et al. (Zerbi et al., 2015) as our reference atlas. This atlas provides a standardized parcellation of the mouse brain into six broadly recognized macroscopic functional systems that have been consistently found in multi-center study (Grandjean et al., 2020): somatosensory, sensorimotor (motor, visual, auditory), olfactory, limbic, basal ganglia, and cerebellum.

##### Bidirectional spatial mapping: Specificity and Coverage

To quantify the structural correspondence between our inferred communities and the canonical RSNs, we computed two complementary geometric metrics: *Specificity* and *Coverage*. Specificity measures the proportion of an inferred community’s spatial volume that falls strictly within a single canonical RSN. Reciprocally, Coverage measures the proportion of a canonical RSN’s spatial volume captured by an inferred community. These metrics were computed for each organizational scheme separately and averaged (weighted by *ω_k_*) to yield an expected spatial mapping per bootstrap resample (see 5.1.10 for mathematical formulations).

##### Quantifying integration: Effective network mappings

To translate these spatial overlaps into interpretable topological principles (i.e., segregation versus integration), we utilized an information theoretic approach. By treating a community’s specificity profile across the six canonical systems as a probability distribution, we calculated its Shannon entropy to derive the *effective number* of spanned RSNs (see 5.1.10 for mathematical formulations). A value near 1 indicates a highly segregated community nested within a single canonical system, whereas a value greater than 1 quantitatively identifies an integrative community bridging multiple systems. We applied this identical entropy framework symmetrically to the coverage profiles to determine whether a canonical system remains cohesive or is functionally subdivided by our generative framework.

#### 4.3.5 Characterizing structural variation within individual communities

An organizational scheme represents a distinct macroscopic topological configuration of the functional connectome. Across the probability landscape of different schemes, a specific community may exhibit varied spatial boundaries, or merge with another community or split. Here we zoom in and characterize the topological multiplicity of the individual communities themselves.

##### Discovering distinct topological patterns: generative clustering

For a given community, we extracted its representative membership probability vectors (the *c*-th column of the *π* matrix) from every organizational scheme across all 500 bootstrap resamples. Crucially, these vectors were strictly masked to include only the community’s surviving ROIs (as defined in 4.3.3). This masking is a necessary prerequisite: high-dimensional clustering algorithms are extremely sensitive to uninformative dimensions (Raftery and Dean, 2006; Bouveyron and Brunet-Saumard, 2014). Including regions that primarily contribute background noise obscures true spatial boundaries, causing the algorithm to overfit and artificially divide the data.

We grouped these high-signal vectors using Gaussian Mixture Models (GMMs) (Murphy, 2012). We utilized GMMs because, similar to the multimodal mixture model used in Section 4.3.2, they are probabilistic generative models. By modeling the ensemble as a mixture of Gaussians, the algorithm identifies *K* dominant centers of mass within the solution space. For each cluster *k*, the GMM yields two key outputs: (1) The mixing weight (*ω_k_*), representing the relative prominence of the cluster in the ensemble, and (2) The centroid (*v*^(^*^k^*^)^), a node-membership vector defining the exact topological structure (i.e., the distinct spatial pattern) represented by the cluster.

Unlike our previous non-parametric models, GMMs require the number of clusters to be provided *a priori*. To objectively determine the optimal number of spatial patterns, we evaluated candidate GMMs using the Bayesian Information Criterion (BIC). Consistent with the information-theoretic principles used for our macroscopic model selection, the BIC can be interpreted as the total number of bits required to compactly encode the ensemble of membership vectors using *k* Gaussian clusters. It resolves the statistical tradeoff between accuracy and complexity by rewarding goodness-of-fit while strictly penalizing the addition of unnecessary parameters (see 5.1.11 for algorithmic details). The minimum BIC value thus identifies the “sweet spot” of the data, revealing the most parsimonious number of spatial patterns statistically supported by the ensemble. Crucially, this data-driven optimization does not force multiplicity; if a community exhibits no meaningful structural variation across the landscape, the minimum BIC will correspond to a single cluster (*k* = 1).

##### Topological merging of redundant patterns

While the minimum BIC identifies the statistically optimal number of Gaussian components, generative mixture models optimize strictly for probabilistic likelihood and do not inherently encode topological constraints (Theis et al., 2016). Consequently, a GMM may artificially split a single continuous spatial pattern into multiple mathematically distinct components simply due to magnitude differences (e.g., one cluster having uniformly stronger or weaker membership probabilities overall) (Hennig, 2010).

Because resolving this redundancy cannot be done using likelihood alone, we consolidated clusters based on a neurobiological criterion: spatial collinearity. We first applied the community’s survival mask (defined in 4.3.3) to the GMM centroids, zeroing-out regions whose expected membership was statistically zero. We then measured the shape redundancy of these masked centroids using cosine distance, a geometric metric that evaluates spatial alignment while perfectly ignoring magnitude differences. Clusters whose masked centroids had a cosine distance less than expected by chance (assessed via a spatially shuffled null model, FDR-corrected; see 5.1.11) were formally merged. The final merged clusters defined the distinct patterns in which the community manifests, and their weights (*ω_k_*) were summed to reflect the total prominence of the distinct pattern.

##### Isolating structurally stable and structurally variable ROIs

Having identified the distinct ROI assignment patterns a community exhibits across the landscape, our final objective was to pinpoint exactly which ROIs drive this structural variation. We evaluated the assignment behavior of individual ROIs by comparing their distributions of assignment probabilities across the distinct patterns using a Mann-Whitney U or Kruskal-Wallis H-test (a non-parametric one-way ANOVA). We tested the null hypothesis that an ROI’s assignment distribution is identical across all patterns. To ensure statistical validity, we first corrected for intra-sample dependency—arising because a single bootstrap resample can yield multiple organizational schemes, and thus multiple dependent assignment vectors—by calculating the median of those dependent vectors (FDR-corrected, *α* = 0.05; see 5.1.11 for exact statistical procedures).

ROIs whose assignment probabilities exhibited no statistically significant difference across the patterns were classified as *structurally stable* ROIs — regions whose functional affiliation remains immutable. Conversely, ROIs exhibiting significant cross-pattern variance were classified as *structurally variable* ROIs — the boundary regions responsible for the community’s structural variation.

## 5 Supplementary

### 5.1 Methods

#### 5.1.1 Functional network aggregation

To ensure statistical validity when aggregating functional connectivity matrices across hierarchical levels (runs and subjects), we utilized Fisher’s *r*-to-*z* transformations (Zar, 1999). Raw Pearson correlation matrices (*r*) were first converted to *z*-scores to normalize the variance of the correlation coefficients.

To account for variations in the amount of usable data caused by frame-scrubbing during pre-processing, the *z*-scored matrices were combined using a weighted average. Each run’s matrix was weighted by its available degrees of freedom (i.e., the total number of valid, un-scrubbed fMRI time-points) (Power et al., 2014). This identical weighted-averaging procedure was repeated to aggregate the subject-level *z*-matrices into the final cohort-level matrix. Finally, the group-average *z*-scored matrix was converted back to Pearson *r* correlation coefficients using the inverse Fisher transformation prior to proportional thresholding.

#### 5.1.2 Mathematical formulation of the Bayesian framework

Our approach is grounded in statistical inference, which allows us to determine the probability that a given latent community structure explains the observed functional network data. This involves defining a forward generative model and subsequently inverting it using Bayesian inference.

##### Generative model

We begin by defining a forward process for how a network *A* (an *N* × *N* adjacency matrix) is constructed. The process assumes that the network’s structure is governed by a latent community partition, *b* = (*b*_1_*,…,b_N_*), which assigns each of the *N* nodes to one of *B* communities. The specific placement of edges between these communities is controlled by a set of parameters, *θ*.

The entire generative process is captured by a joint probability distribution:

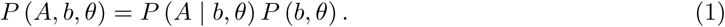

Here, the likelihood *P* (*A* | *b, θ*) specifies the probability of generating the observed network *A* given a specific partition and parameter set. The prior *P* (*b, θ*) encodes our structural assumptions about the network’s organization (e.g., hierarchical or assortative) before any data is observed.

##### Non-parametric generative procedure

We utilize a non-parametric version of this framework, meaning the number of communities (*B*) is inferred directly from the data rather than being fixed *a priori*. The forward generative procedure is inherently sequential: we first sample the number of communities *B*, then their relative sizes, then the specific partition *b* (assigning individual nodes to communities), and finally the edge parameters *θ* from the prior distribution. Edges are then placed between communities according to the likelihood distribution to construct the final network.

##### Bayesian inference and the Minimum Description Length

Given the empirically observed functional network *A*, we invert the generative process using Bayes’ rule to infer the posterior distribution of plausible partitions:

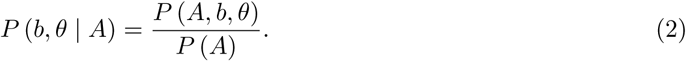

The denominator is the model evidence: the total probability that the generative model assigns to the observed network, integrated over all possible partitions.

To operationalize this search, the joint probability can be reframed from an information-theoretic standpoint by converting it into the description length, Σ:

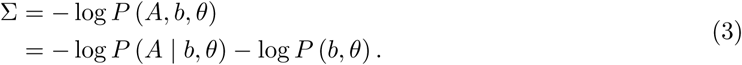

The description length represents the total number of bits required to describe the network’s structure using the chosen model. It consists of the bits needed to describe the data given the model (the negative log-likelihood) and the bits needed to describe the model parameters themselves (the negative log-prior). According to the Minimum Description Length (MDL) principle, maximizing the posterior probability is mathematically equivalent to finding the partition *b* that minimizes Σ.

#### 5.1.3 Mathematical formulations of generative models

The mathematical form of the Stochastic Block Model (SBM) is derived directly from the principle of maximum entropy (Jaynes, 2003). The SBM is the most principled, maximally random generative model subject to a single constraint: the expected number of edges between nodes in any two blocks *r* and *s*. In the sparse limit typical of large empirical networks, the probability of observing multiple edges between two nodes is negligible, allowing the exact likelihood to be approximated by a Poisson distribution (Peixoto, 2019).

##### The non-degree-corrected standard SBM (nd-s-SBM)

This foundational model (Peixoto, 2017) assumes nodes in the same block are statistically equivalent. Using a Poisson formulation, the likelihood of observing a network *A* given a partition *b* and a matrix of expected edge counts *λ* is:

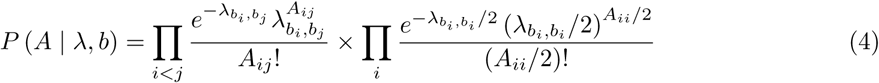

Here, *λ_r,s_*is the expected number of edges between block *r* and block *s*. The prior distributions are explicitly defined to prevent biasing the model complexity:

- **Prior on the partition,** *P* (*b*): We use a non-parametric hierarchical prior that sequentially samples the number of blocks *B*, the block sizes, and the partition *b* from uniform distributions.
- **Prior on edge counts,** *P* (*λ*): We assume edge counts between blocks are completely random, sampled from a uniform distribution.

##### The degree-corrected SBM (dc-s-SBM)

To capture heterogeneous degree distributions not determined by community structure, the dc-s-SBM (Peixoto, 2017) introduces a degree parameter, *θ_i_*, for each node. The modified likelihood is:

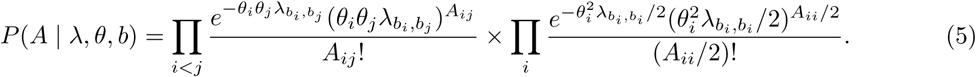

##### Hierarchical and Overlapping SBMs

The hierarchical SBM (h-SBM) places a hierarchical prior over the edge counts *λ*, positing that the connection matrix is structured and described by another SBM at a higher level (Peixoto, 2014c). The overlapping SBM (o-SBM) shifts the assignments from nodes to half-edges, creating a mixture model where a node’s fraction of edges in each community dictates its mixed-membership proportions (Peixoto, 2015). Both models can be parameterized with or without degree-correction (*θ_i_*).

#### 5.1.4 The assortative benchmark and modularity

Modularity maximization (Newman, 2006) aims to find a partition *b* by optimizing a quality function, *Q*, with a resolution parameter, *γ*:

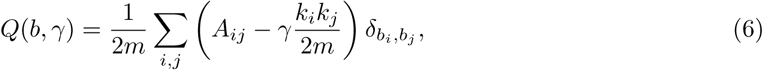

where *m* is the total number of edges, *k_i_* is the degree of node *i*, and *δ_b__i,bj_* = 1 if nodes *i* and *j* are in the same community. This descriptive method exclusively rewards assortative communities.

Its generative equivalent, the assortative SBM (a-SBM) or Planted Partition model, is a constrained version of the dc-s-SBM. Instead of a full matrix *λ*, its likelihood assumes only two connection rates: a within-group rate (*λ_in_*) and a between-group rate (*λ_out_*). Because maximizing *Q* is mathematically equivalent to maximizing the likelihood of the a-SBM under specific limits (Newman, 2016), comparing the inference results of the a-SBM against the heuristic output of *Q* provides a direct test of whether the assortative structure is statistically genuine or a product of algorithmic overfitting.

#### 5.1.5 Markov chain Monte Carlo implementation details

To infer the community structure, we must find the partitions (*b*) that have the highest posterior probability, *P* (*b* | *A*), corresponding to the lowest description lengths (Σ). Because the space of all possible partitions is too vast to search exhaustively, we approximate the posterior distribution using the advanced Markov chain Monte Carlo (MCMC) algorithms developed by Peixoto (Peixoto, 2014a, 2020).

Standard MCMC algorithms propose moving a single node to a new community chosen uniformly at random. This results in exceedingly slow mixing times and trapping in local minima. The graph-tool implementation overcomes this by utilizing two specialized move proposals:

1. **Neighborhood-guided local moves:** When proposing a new community assignment *r* for node *i*, the target community is not chosen at random. Instead, it is proposed with a probability proportional to the number of existing connections between the neighbors of node *i* and the nodes already residing in community *r* (Peixoto, 2014a). This leverages local network topology to guide the search aggressively toward more probable solutions.
2. **Macroscopic merge-split moves:** To escape deep local minima separated by valleys of low probability, the algorithm periodically proposes merging two entire communities together, or splitting a single community into two (Peixoto, 2020). This allows the Markov chain to make macroscopic leaps across the posterior landscape that would be impossible via single-node updates.

Regardless of the proposal type, all moves are formally evaluated using the Metropolis-Hastings criterion (Metropolis et al., 1953; Hastings, 1970). If a proposed move decreases the description length, it is strictly accepted. If it increases the description length, it is accepted with a probability proportional to the difference in description lengths, ensuring the algorithm maintains detailed balance while exploring the global landscape.

#### 5.1.6 Validating model faithfulness: MCMC diagnostics

To quantitatively validate that a model produced a coherent posterior landscape, we assessed MCMC convergence based on the distributions of the description length (Σ) sampled by each of the five independent chains.

First, we visually compared the posterior distributions (histograms) of Σ utilizing only samples collected during the second half of the chains (ensuring the chains were sufficiently past the initial equilibration period). Substantial overlap provided initial evidence that the chains were sampling from the same underlying landscape.

We then formally quantified this overlap using the Kolmogorov-Smirnov (KS) distance, a non-parametric statistical test that measures the maximum absolute difference between two empirical cumulative distribution functions (CDFs) (Bertsekas and Tsitsiklis, 2008). For two MCMC chains with empirical CDFs *F*_1_(*x*) and *F*_2_(*x*), the KS distance is defined as:

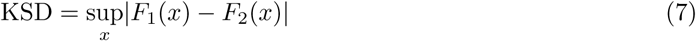

We tracked the KSD for all 10 possible pairwise comparisons between the five independent chains across all MCMC steps. A model was formally considered to have converged only if the maximum distance for all pairwise comparisons fell strictly below a standard statistical threshold of 0.2 (Gibbons and Chakraborti, 2011), indicating no significant macroscopic difference between the sampled posterior distributions.

#### 5.1.7 Model comparison: Total Description Length

After identifying a set of faithful models, we performed a formal model comparison to determine which provided the most parsimonious explanation of the data. The principled Bayesian approach for comparing two models, M_∞_ and M_∈_, is to compute the posterior odds ratio. Assuming no *a priori* preference for either model, this simplifies to the Bayes factor, which is the ratio of their marginal likelihoods (model evidences):

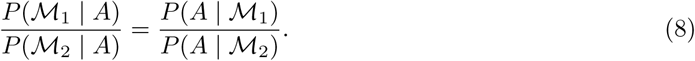

The model evidence, *P* (*A* | M), is the probability of observing the network data *A* given the model M, integrated over all possible partitions. A higher evidence indicates a more probable model.

For practical comparison, we work with the negative logarithm of the evidence, expressed as the Total Description Length (TDL). The evidence can be written as an average over the posterior distribution, *q*(*b*) = *P* (*b* | *A*):

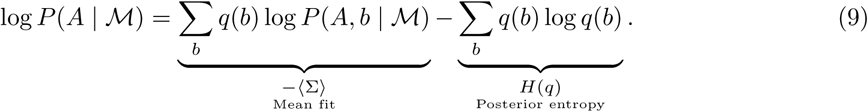

Here, the first term −⟨Σ⟩ is the negative expected description length over the posterior distribution, and the second term *H*(*q*) is the Shannon entropy of the posterior.

The TDL therefore explicitly accounts for both the fit per partition and the structural uncertainty in the solutions:

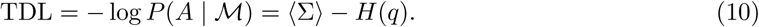

⟨Σ⟩ represents the average goodness-of-fit over all highly probable partitions sampled by the MCMC chains after the equilibration period. *H*(*q*) quantifies the uncertainty arising from the diversity of the solution landscape. By penalizing models with high posterior entropy, the TDL enforces a strict preference for models whose posterior distributions are mathematically compact and well-connected.

Because the posterior entropy *H*(*q*) is intractable to compute directly by exhaustively enumerating all possible partitions, we approximate it using the output of a multimodal mixture model (described in 5.1.8). The entropy is decomposed into two parts: the average entropy *within* each mode, and the entropy *between* the modes (calculated from their transition probabilities). This approach, implemented via the graph-tool Python library (Peixoto, 2014b), yields a robust and principled TDL for direct model adjudication.

#### 5.1.8 Characterizing the posterior distribution

The raw output from the MCMC sampling is a large ensemble of possible partitions. Before this ensemble can be interpreted, it must be rigorously processed to account for label ambiguity, landscape multiplicity, and topological redundancy. All implementations of the random label and mixture models were performed using the graph-tool Python library (Peixoto, 2014b).

##### Aligning the partitions: the random label model

The ensemble of partitions inevitably contains arbitrary group labels for equivalent communities (label switching). To resolve this, we applied the random label model (Peixoto, 2021). Formally, this model posits the existence of a hidden, canonical partition *c*. Each observed MCMC sample *b* is treated as a realization of this canonical partition that has been subjected to an unknown, random label permutation.

To align an observed partition to the canonical template, we must infer the specific permutation that maximizes their structural overlap. This exact inference is computationally framed as a maximum weighted bipartite matching problem. We construct a bipartite graph where the two sets of nodes represent the community labels of *b* and *c*, and the edge weights represent the number of shared nodes (the joint contingency table) between any two communities. The model utilizes the Kuhn-Munkres (Hungarian) algorithm (Kuhn, 1955; Munkres, 1957; Jünger et al., 2010) to efficiently find the exact label assignment that maximizes this bipartite weight. To align an entire ensemble where the canonical template is initially unknown, the model employs an iterative approach: it aligns random partitions to a growing reference set, continuously updating a node co-occurrence matrix to refine the consensus template until the global assignments stabilize.

##### Summarizing the posterior: the multimodal mixture model

To address posterior heterogeneity, we summarized the aligned landscape using a multimodal mixture model (Peixoto, 2021). This clustering algorithm assumes the sampled ensemble of partitions is drawn from a mixture of *K* distinct structural modes. It adaptively determines the optimal number of modes via a greedy optimization procedure that iteratively reassigns partitions, merges similar modes, and splits heterogeneous ones.

Crucially for our downstream analyses, this procedure yields two key parameters that summarize the landscape: the mode probabilities (*ω_k_*, where *_k_ ω_k_* = 1) and the within-mode node-membership probability matrices (*π*^(^*^k^*^)^). This explicitly separates the statistical uncertainty *between* competing global configurations (*ω_k_*) from the topological uncertainty *within* a given mode (*π*^(^*^k^*^)^). For the complete mathematical derivation of this generative mixture process, including the exact marginalization over hidden label permutations, we refer the reader to Peixoto (2021).

##### Topological merging of redundant modes

To isolate neurobiologically distinct organizational schemes, we evaluated the structural redundancy between all pairs of modes. To ensure dimensional consistency prior to comparison, each *N* × *B_k_* node-membership probability matrix (*π*^(^*^k^*^)^) was first zero-padded to a standardized size of *N* × *B_max_*, where *B_max_* is the maximum number of communities observed across all *K* modes. These standardized matrices were then flattened into one-dimensional vectors, allowing us to calculate the pairwise cosine similarity between all modes. To test whether a high similarity score indicated true topological redundancy rather than incidental structural overlap, we generated a spatial null distribution by randomly shuffling the ROI indices 1,000 times and recalculating the similarity.

An empirical *p*-value was derived based on the frequency with which the null similarity exceeded the true observed similarity. To rigorously control the family-wise error rate across all pairwise mode comparisons, we applied Benjamini-Hochberg False Discovery Rate (FDR) correction (*α* = 0.05) (Benjamini and Hochberg, 1995). Using these FDR-corrected *p*-values, we constructed a binary adjacency graph where nodes represented individual mixture modes, and edges connected pairs exhibiting statistically significant topological redundancy. Finally, we extracted the connected components of this graph to formally merge the modes. The probabilities (*ω_k_*) of all constituent modes within a connected component were summed to yield the total probability of the newly consolidated organizational scheme, and their zero-padded node-membership matrices (*π*^(^*^k^*^)^) were combined via a weighted average (proportional to *ω_k_*) to define its consensus topological structure.

##### Aligning schemes across bootstrap resamples

To map the organizational schemes identified in each bootstrap resample back to the reference schemes, we computed a pairwise cosine distance matrix between their respective flattened *π* matrices. We formally treated this mapping as a linear sum assignment problem and solved it using the Hungarian algorithm (Kuhn, 1955; Munkres, 1957; Jünger et al., 2010) to establish an optimal one-to-one bipartite matching. Because the number of distinct schemes may naturally differ between a bootstrap resample and the reference ensemble, this assignment was computed on a rectangular distance matrix, allowing the algorithm to match the maximum possible pairs while leaving any excess, unmatchable schemes unassigned.

Crucially, an assignment algorithm will force a match for these paired subsets even if their absolute distance is large. To ensure these represented genuine topological matches rather than forced algorithmic pairings, we subjected the optimally assigned matrix pairs to the exact same spatial permutation test described above (1,000 ROI shuffles, FDR-corrected *α* = 0.05). Matches that failed to exceed this spatial null threshold were discarded. This rigorous mapping and validation allowed us to reliably track specific organizational schemes, and their probabilities (*ω*), across the population variance.

#### 5.1.9 Community survival analysis

To rigorously define the spatial boundaries of the inferred communities, we isolated the reliable expected membership of each ROI. For a given bootstrap resample, the expected node-membership matrix, E[*π*], was calculated as the weighted average of the scheme-specific matrices (*π*^(^*^k^*^)^), weighted by their respective probability of occurrence (*ω_k_*):

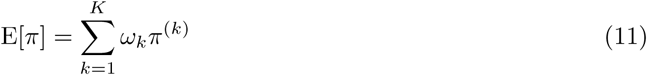

where *K* is the total number of distinct organizational schemes identified in that specific resample.

Next, we compiled the distribution of these expected membership values for each ROI across the 500 bootstrap resamples to test the null hypothesis that an ROI’s true expected membership in a given community was zero. For every element in the expected node-membership matrix (that is, the probability of a specific ROI belonging to a specific community), an empirical *p*-value was calculated. This *p*-value was defined as the proportion of the bootstrap distribution that was equal to zero.

To rigorously control the false discovery rate across the entire membership matrix (which involves simultaneous tests for all ROIs across all identified communities), we pooled all empirical *p*-values and applied a single Benjamini-Hochberg False Discovery Rate (FDR) correction (*α* = 0.05) (Benjamini and Hochberg, 1995). Matrix elements with an FDR-corrected *p*-value *<* 0.05 (conceptually equivalent to a zero-excluding, globally adjusted confidence interval) were deemed to have “survived” the thresholding process. The ROIs corresponding to these surviving elements formed the definitive binary mask for each community, establishing the feature space used in subsequent analyses.

#### 5.1.10 Resemblance to canonical systems: spatial mappings and effective numbers

To quantitatively assess the alignment between our inferred communities and the canonical RSNs, we represented both as voxel-wise brain volumes to account for varying ROI sizes. Each of the *R* canonical RSNs was represented as a binary volume, *w^r^* ∈ {0, 1}*^Nv^*, where *N_v_* is the total number of voxels.

For a given organizational scheme *k*, an inferred community *c* is defined by its node-membership probability vector (the *c^th^*column of *π*^(^*^k^*^)^). We constructed a continuous spatial volume for the community, *v^c^*∈ [0, 1]*^Nv^*, by broadcasting these node-wise probabilities to all voxels within their corresponding ROIs.

We quantified spatial alignment using a bidirectional analysis per organizational scheme *k*:

1. **Specificity (***S*): From the community’s perspective, specificity (*S*^(^*^k^*^)^) was calculated as the fraction of community *c*’s total volume that lies within RSN *r*:

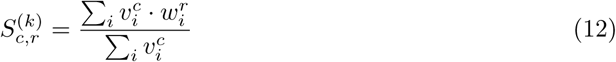

where *i* indexes voxels.

1. **Coverage (***C*): From the RSN’s perspective, coverage (*C*^(^*^k^*^)^) was calculated as the fraction of RSN *r*’s total volume covered by community *c*:

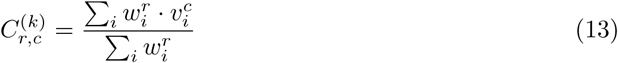

For a given bootstrap resample, these scheme-specific matrices (*S*^(*k*)^ and *C*^(*k*)^) were combined into expected mapping matrices by taking a weighted average across all identified schemes, weighted by their respective probability of occurrence (*ω_k_*).

To measure how broadly a community is distributed across systems, we calculated the Shannon entropy of its expected specificity profile (its row in the expected *S* matrix) and exponentiated the result to yield the *effective number* of RSNs community *c* spanned:

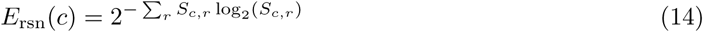

This identical entropy derivation was applied symmetrically to the rows of the expected coverage matrix (*C*) to calculate the effective number of communities that covered a canonical RSN *r* (*E*_comm_(*r*)).

#### 5.1.11 Characterizing structural multiplicity of individual communities

##### Dimensionality reduction for generative clustering

Prior to clustering, the membership vectors were strictly masked to include only the surviving ROIs of the community. This masking is a mathematical necessity for density-based generative models like GMMs. In high-dimensional spaces, the inclusion of dimensions (ROIs) that consist primarily of background noise fundamentally alters the geometry of the data distribution. Because the volume of the space increases exponentially with dimensionality, data points become sparse, and the distance (or density) contrast between true clusters and noise diminishes—a phenomenon known as the curse of dimensionality (Bouveyron and Brunet-Saumard, 2014). By masking out non-surviving ROIs, we restrict the GMM to a high-signal subspace, allowing it to accurately model the covariance structure and estimate the true likelihoods of the underlying topological patterns.

##### Model selection via Bayesian Information Criterion

All GMM fitting and evaluation procedures were implemented using the scikit-learn library in Python (Pedregosa et al., 2011). For a given community, GMMs were fitted to the masked membership vectors across a range of candidate cluster counts (*k* = 1, 2, · · ·, 10) using maximum likelihood estimation. To objectively identify the optimal number of spatial patterns, we selected the model that minimized the Bayesian Information Criterion (BIC). The BIC formalizes the tradeoff between a model’s goodness-of-fit and its complexity:

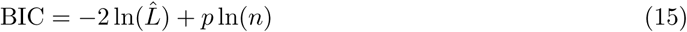

where *L*^^^ is the maximized likelihood of the model given the data, *p* is the total number of estimated parameters (which increases with *k*), and *n* is the number of membership vectors. The first term rewards models that accurately capture the data’s structure, while the second term strictly penalizes the addition of unnecessary Gaussian components, thereby preventing overfitting.

##### Topological merging of redundant patterns

To resolve structural redundancy among the optimal GMM components, we evaluated the spatial collinearity of their centroids (mean vectors, *v*^(^*^k^*^)^) using cosine distance. For two centroids *i* and *j*, the cosine distance is:

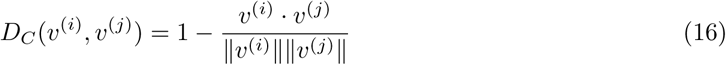

This metric isolates the relative distribution of membership strengths across ROIs (the shape) while ignoring uniform differences in probability magnitude.

To determine statistical significance, we generated a spatial null distribution by randomly shuffling the ROI indices of the centroids 1,000 times and recalculating the pairwise cosine distances. An empirical *p*-value was calculated for each pair based on the proportion of null distances that were smaller than the true observed distance. We pooled these *p*-values and applied Benjamini-Hochberg False Discovery Rate (FDR) correction (*α* = 0.05) (Benjamini and Hochberg, 1995). Clusters exhibiting statistically significant shape redundancy were merged via connected components. The centroid of each newly merged entity served as the representative spatial pattern, and the mixing weights (*ω_k_*) of its constituent GMM clusters were summed.

##### Statistical isolation of stable and flexible ROIs

To rigorously execute the Kruskal-Wallis H-test for individual ROIs across the distinct patterns, we first had to ensure the observations were statistically independent. Because a single bootstrap resample can yield multiple organizational schemes, it can contribute multiple membership vectors to the overall ensemble. Testing these vectors directly would violate the assumption of independence and artificially inflate the degrees of freedom.

To correct for this intra-sample dependency, we calculated the median of all membership vectors that originated from the same bootstrap resample *and* were assigned to the same distinct spatial pattern. This compression yielded a distribution of independent expected membership values for each pattern (with a maximum of 500 independent observations per pattern). Using these independent distributions, we performed the Kruskal-Wallis H-test for each ROI. Finally, we applied FDR correction (*α* = 0.05) across the tests for all surviving ROIs to classify them as either stable (failing to reject the null hypothesis) or flexible (exhibiting significant cross-pattern variance).

### 5.2 Communities at the ground and upper level of the hierarchy

**Figure 11:**
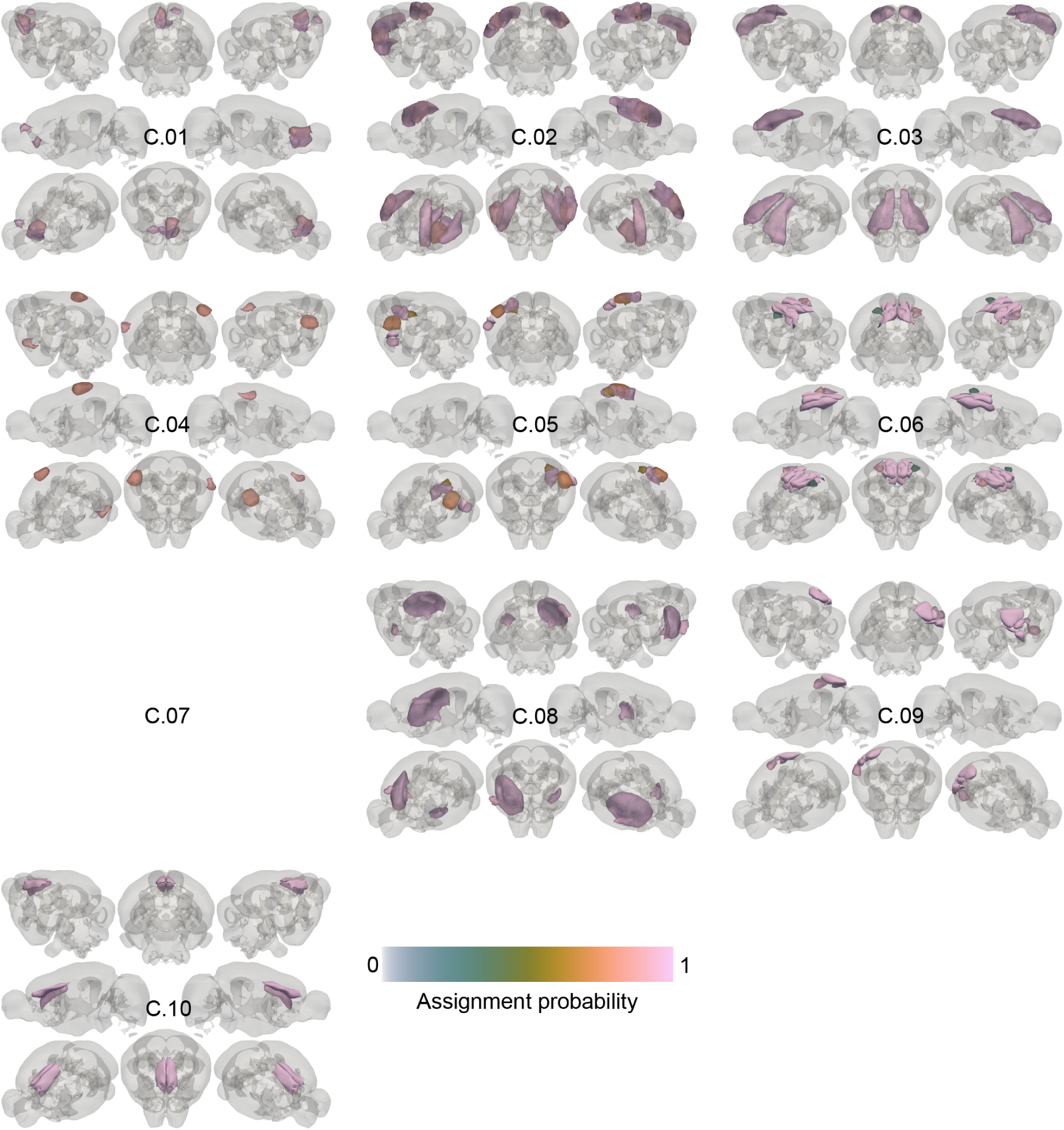
First 10 communities at the ground level that merge into 9 communities at the middle level.

**Figure 12:**
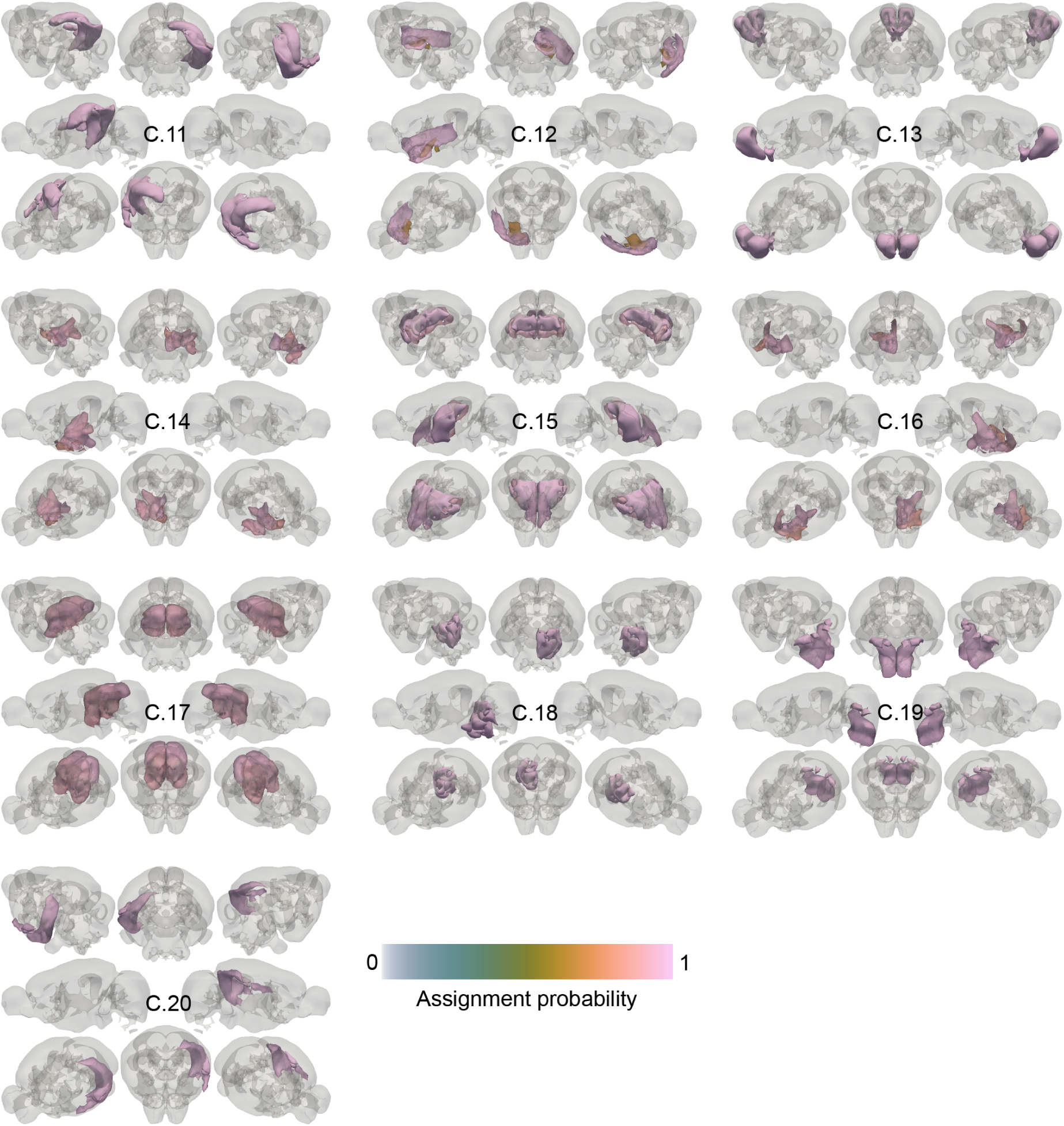
Second 10 communities at the ground level that merge into 9 communities at the middle level.

**Figure 13:**
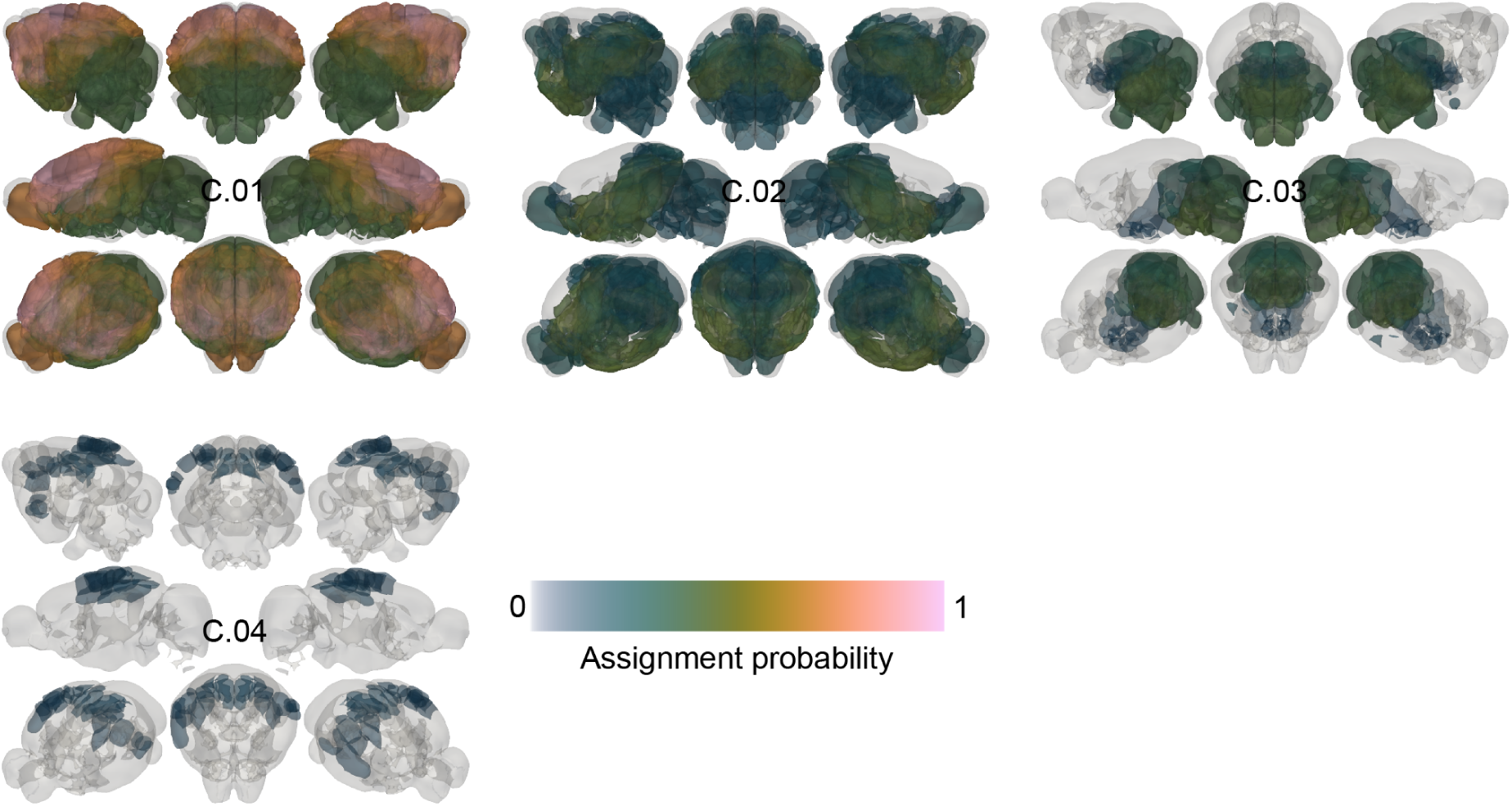
Communities at the ground level that merge into 9 communities at the middle level.

### 5.3 Multiplicity in organizational schemes for the modularity maximization method

To verify that the multiple organizational schemes observed in our results are a genuine property of the functional connectome under a generative framework, rather than an algorithmic artifact of our multi-modal mixture model pipeline, we applied the exact same Bayesian inference procedure to partitions generated via traditional modularity maximization Newman and Girvan (2004).

The modularity objective landscape is known to be a highly irregular “rugged plateau” of structurally similar partitions Good et al. (2010). If our clustering algorithm artificially forces continuous variation into distinct macroscopic modes, we would expect to see false multiplicity here as well.

Instead, applying our clustering pipeline to the modularity maximization results yielded an effectively unimodal landscape. While our procedure detected two schemes, their occurrence probabilities were massively skewed (Figure 14). A single dominant organizational scheme occurred almost exclusively (*ω*_1_≈ 1), while the alternative scheme was exceptionally rare (*ω*_2_ ≈ 0). Nonparametric statistical testing confirmed this profound asymmetry in prevalence (large effect size, *p <* 0.05). Furthermore, the topological differences between this dominant scheme and the rare variant were exclusively localized to minor reconfigurations in the hindbrain (not shown); the entire cortical and subcortical functional architecture remained invariant across the landscape. Because the clustering algorithm successfully collapsed the rugged modularity plateau into a single unified cortico-subcortical architecture, this suggests that our procedure does not artificially hallucinate multiplicity. Consequently, the distinct, co-dominant schemes identified by our generative framework may reflect genuinely distinct whole-brain functional organizations.

**Figure 14:**
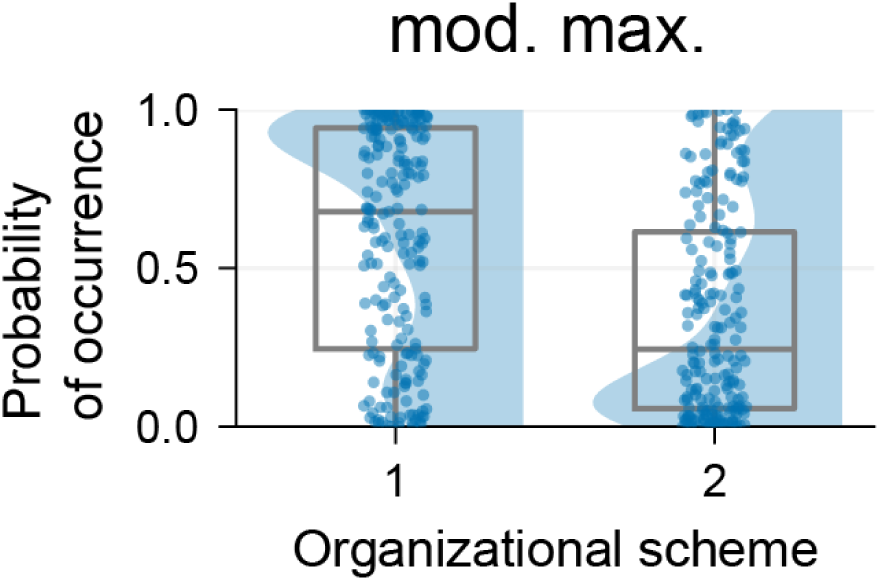
Modularity maximization yields an effectively unimodal solution landscape. Occurrence probabilities (*ω*) of the two organizational schemes detected by applying the MCMC + mixture model procedure to partitions generated via modularity maximization. Unlike the distinct and co-dominant schemes identified by the generative framework, modularity maximization results in a single dominant scheme (*ω*_1_ ≈ 1; mode of the distribution) and a rare alternative (*ω*_2_ ≈ 0). This profound asymmetry suggests that our proposed framework effectively absorbs minor topological fluctuations into a unified scheme and does not artificially inflate structural variation.

### 5.4 Leave-one-out (LOO) analysis for analyzing source of multiple organizational schemes

To verify that the observed multiplicity in organizational schemes reflects a fundamental property of the functional connectome rather than an artifact driven by an outlier subject, we evaluated the robustness of the multi-modal landscape against single-subject omission. We conducted a systematic leave-one-out (LOO) analysis, inferring the posterior landscape for independent (*n* − 1)-sized cohorts. As detailed in Table 1, multiple organizational schemes (≥ 2) were successfully recovered in every LOO iteration. This suggests that multiplicity is a generalizable property of the resting-state architecture rather than an outsized influence of a single subject.

**Table 1:** Robustness of structural multiplicity to single-subject omission. A leave-one-out (LOO) analysis suggests that the multi-modal posterior landscape is not driven by any single outlier subject. Every (*n* − 1)-sized cohort consistently yielded multiple organizational schemes.

| Omitted Subject | Number of Organizational Schemes |
| --- | --- |
| sub-SLC01 | 3 |
| sub-SLC03 | 5 |
| sub-SLC04 | 2 |
| sub-SLC05 | 3 |
| sub-SLC06 | 7 |
| sub-SLC07 | 2 |
| sub-SLC08 | 4 |
| sub-SLC09 | 4 |
| sub-SLC10 | 3 |

### 5.5 Analyses for 10% network sparsity

We evaluated our Bayesian generative modeling framework for graphs at a sparser edge density (10% edge retention; see Section 4.1.5 for details) to ensure our framework generalizes beyond a specific threshold.

#### 5.5.1 Generative model selection

We computed the Total Description Lengths (TDL) for all the seven organizational principles and observed that the non-degree-corrected hierarchical principle yielded the lowest TDL and the purely assortative model yielded the highest TDL (Figure 15). Additionally, the non-degree-corrected models were consistently more compatible with the data than their degree-corrected variants.

**Figure 15:**
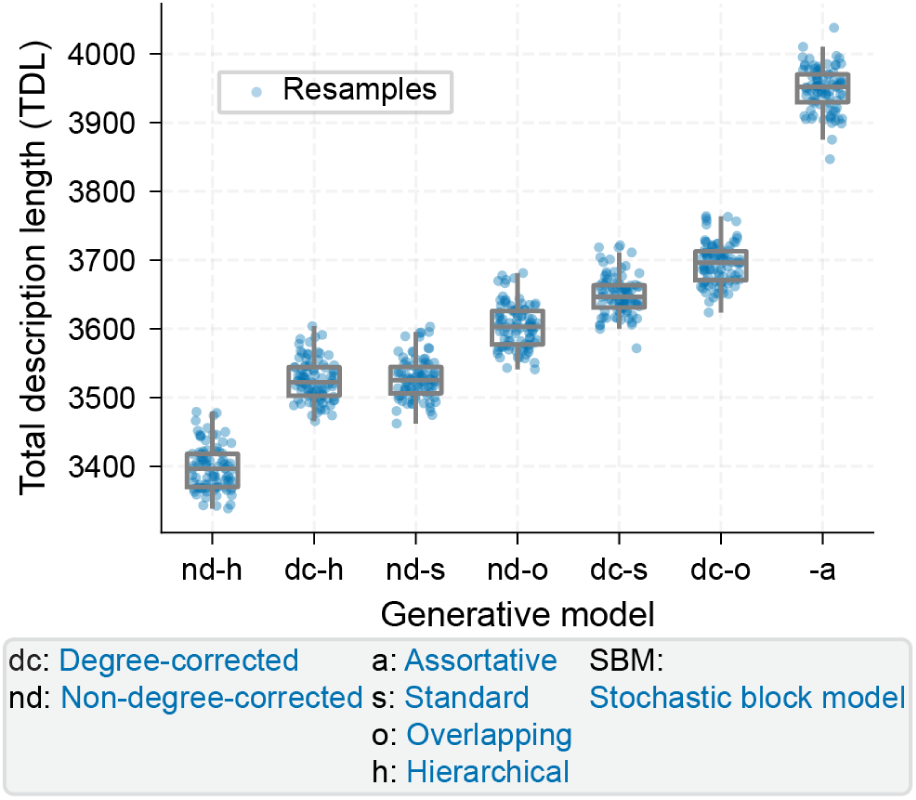
Generative model preferences are robust to hieghtened graph sparsity. Model comparison based on Total Description Length (TDL) across seven competing hypotheses, evaluated at 10% edge density. Consistent with the primary analysis (Figure 4), the non-degree-corrected hierarchical model (nd-h-SBM) provides the optimal fit to the data, while the purely assortative model (a-SBM) remains the least compatible. The uniform preference for non-degree-corrected models (nd) over degree-corrected models (dc) is also preserved.

#### 5.5.2 Inferred communities

We computed the expected community compositions inferred by the optimal non-degree-corrected hierarchical model. The analysis revealed seven communities (Figure 16).

**Figure 16:**
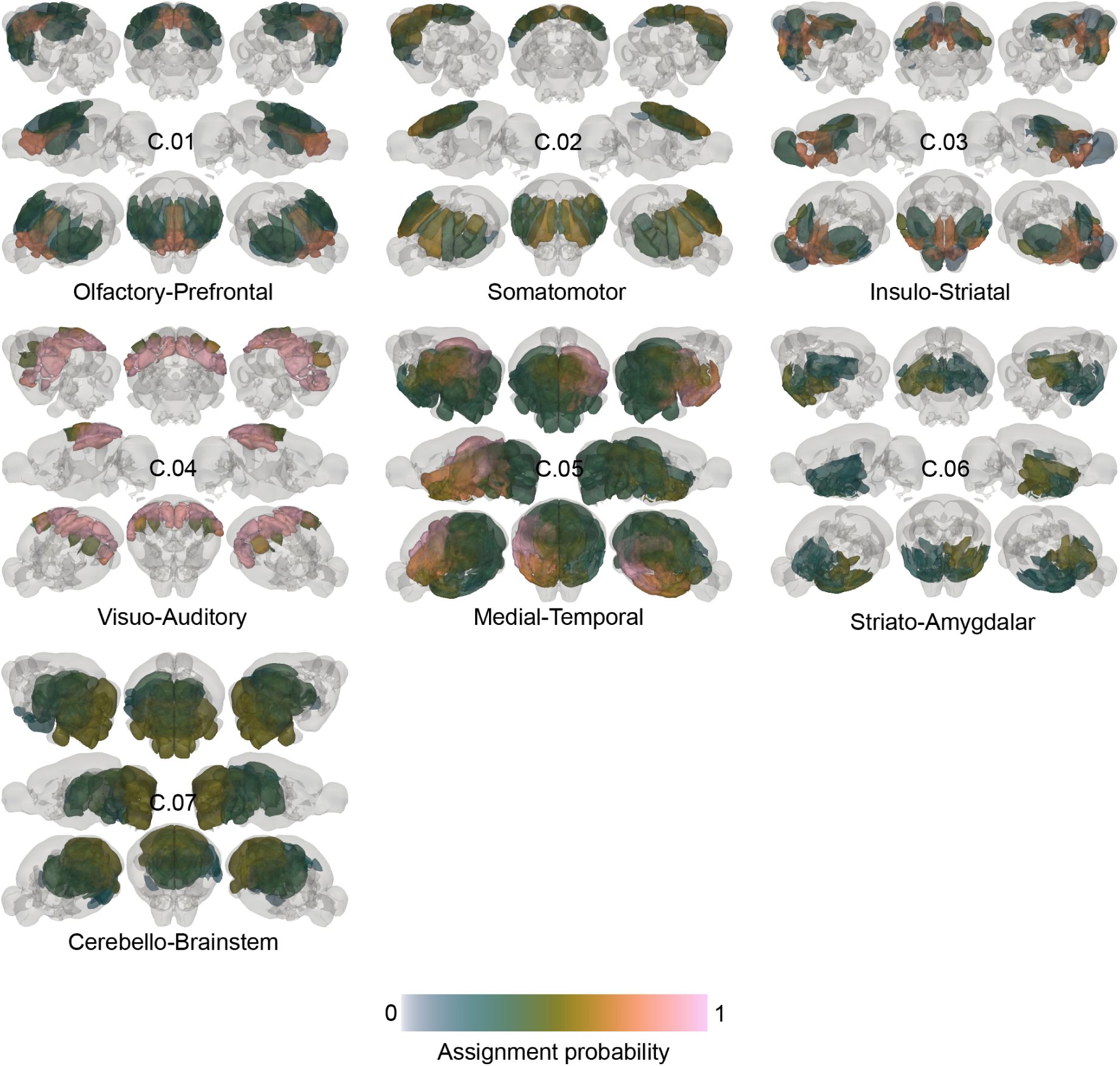
The macroscopic functional architecture condenses parsimoniously at heightened graph sparsity. Expected ROI compositions of the seven middle-level communities inferred by the non-degree-corrected hierarchical model (nd-h-SBM) evaluated at 10% edge density. Due to the removal of weaker edges, the generative model parsimoniously condenses the functional architecture into seven broader macro-systems. Core anatomical structures, such as the Somatomotor and Medial-Temporal communities, remain highly conserved, while tightly related physiological and associative axes fuse into cohesive larger-scale domains.

Compared to the top 20% edge density, the core anatomical axes remained highly stable. The So-matomotor and Medial-Temporal communities were perfectly conserved. Tightly related physiological structures, such as the Cerebello-Brainstem and Ponto-Midbrain, fused into a unified deep-brain axis, while the prefrontal and olfactory architectures similarly condensed.

